# Evolving stem cell fate capacities and transcriptional priming in the developing olfactory epithelium

**DOI:** 10.64898/2026.07.28.741343

**Authors:** Dana Bakalar, Catie Kaneshiro, Chloe Zhao, Tyson Fang, Sandrine Dudoit, Elizabeth Purdom, Kelly Street, John Ngai, Whitney Heavner

## Abstract

The olfactory epithelium of adult mammals contains two populations of stem cells that support its remarkable ability to regenerate neuronal and non-neuronal cell types throughout life. How olfactory epithelial cell types are established during development, however, is not well understood. Here, we use genetic lineage tracing and single-cell RNA sequencing of the perinatal mouse olfactory epithelium to construct a developmental trajectory consisting of multiple lineages. We identify transitional states and lineage relationships between individual cells and establish *Ascl1^+^* cells as the primary multipotent progenitors in the perinatal olfactory epithelium. Further, *Ascl1^+^* cells become progressively restricted in their cell fate capacity over developmental time and appear to be transcriptionally primed toward specific lineages. We also predict signaling pathways that may contribute to lineage plasticity and niche permissiveness. Together, these results contribute to our understanding of how cell-intrinsic and -extrinsic signals contribute to the establishment of a stem cell niche.

## Introduction

Neural tissues consist of many subtypes of specialized neuronal and non-neuronal cells that together drive function. A recurring strategy used to generate cellular diversity in the nervous system is the sequential generation of specific cell types by a single progenitor population (Martynoga et al. 2012). One classic model of cell type diversification, supported by heterochronic transplantation experiments *in vivo* and reaggregation experiments *in vitro* in the vertebrate retina, posits that the competence of progenitor cells to respond to environmental cues changes over time in a stereotyped, sequential manner (Cepko et al. 1996).

The main olfactory epithelium (OE) is an experimentally facile model for investigating interactions between progenitor cells and other cells in the niche. A relatively simple pseudo-stratified epithelium containing neurons and non-neuronal cells, the OE retains extraordinary proliferative and regenerative capacity from embryogenesis through adulthood (Suzuki and Takeda 1993). The adult OE contains two progenitor populations: globose basal cells (GBCs), which are proliferative neural stem cells that sustain ongoing neurogenesis, and horizontal basal cells (HBCs), which are reserve stem cells capable of regenerating all OE cell types after severe injury, including olfactory sensory neurons (OSNs), sustentacular (Sus) cells, cells of the Bowman’s gland (BG), and microvillus cells (MV). Although the lineages arising from HBCs during homeostasis and after injury have been well characterized (Fletcher et al. 2017; Gadye et al. 2017), less is known about how HBCs and GBCs are established, the extent of their respective contributions to OE development, and how much the local microenvironment, including signaling between daughter cells and progenitor cells, affects cell fate.

As early as embryonic day 10.5, a subset of *Sox2+* neural stem cells in the olfactory pit -- the presumptive precursors of the OE (Scriven 1993; Engel et al. 2016) -- begin to express the proneural bHLH transcription factor *Ascl1* (Kawauchi et al. 2005), which is required for neuronal progenitor maintenance (Cau et al. 2002; Krolewski et al. 2012; Murray et al. 2003). Consistent with neural progenitor cells in other tissues, *Ascl1^+^* cells in the OE predominantly generate neurons (Cau et al. 1997, 2000, 2002), but starting from embryonic day 12.5 and throughout OE neurogenesis (Beites et al. 2005), also produce Sus cells (Tucker et al. 2010; Murray et al. 2003; Chen et al. 2014; Gokoffski et al. 2011), MV cells (Yamaguchi et al. 2014), and BG/duct cells (Chen et al. 2014). *Ascl1* may play a role in the development of HBCs, although the lineage relationship is still uncertain (Krolewski et al. 2012; Packard et al. 2011). These studies suggest that the developmental capacity of *Ascl1^+^* cells in the OE may change over time, but the sequence and proportions of cell types generated by *Ascl1^+^*cells have not been systematically examined across development, and the pathways driving these cell fate decisions are still unknown.

In the present study, we use genetic lineage tracing of stem cell populations in the developing OE to identify changes in cell fate capacity and stem cell plasticity over time. We also incorporate single-cell transcriptomics to establish a trajectory of OE development that captures transitional states, lineage relationships between single cells, and changes in gene expression along each lineage. Finally, we identify potential signaling interactions between cell types that may contribute to progenitor capacity and plasticity within the developing niche.

## Results

### Lineage tracing reveals a shift in cell fate capacity of Ascl1^+^ cells

*Ascl1*-expressing cells in the embryonic OE have the potential to generate all sensory OE cell types (Paronett et al. 2023), but less is known about the capacity of these cells during postnatal development. To determine if and how the fate capacity of *Ascl1*^+^ cells changes throughout development, we genetically lineage traced *Ascl1*⁺ cells from several developmental time points in the mouse. Tamoxifen was administered to *Ascl1^CreER/+^*; *R26^tdTomato^* timed-pregnant dams on embryonic day (E) 14.5, or pups on postnatal day (P) 0, 3, 5, or 8, or adults on P35. Embryos were dissected at E19.5, pups were dissected at weaning, and adults were dissected at P49 (**Fig. 1A**).

**Figure 1.**
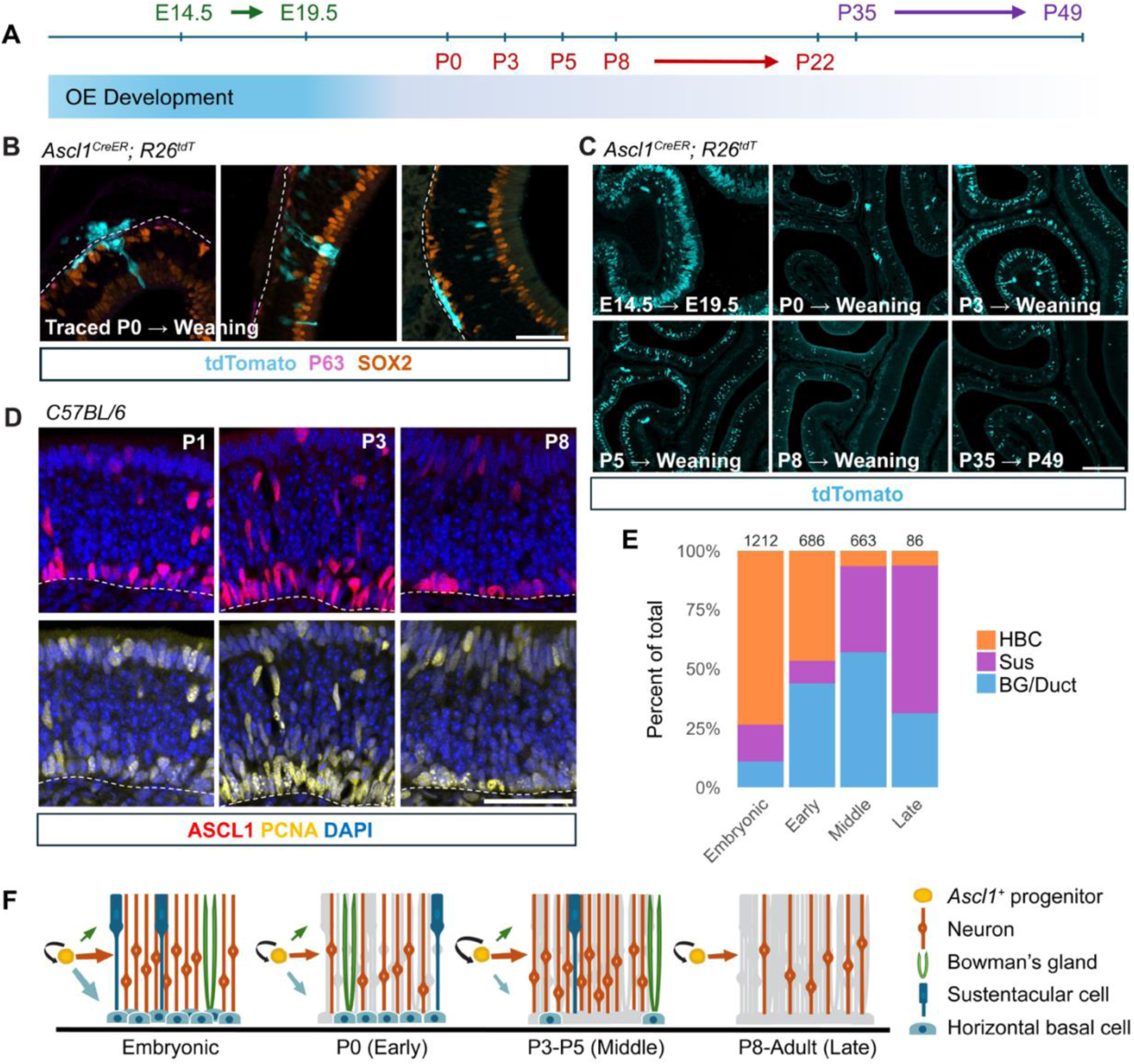
The capacity of *Ascl1*^+^ cells to produce non-neuronal olfactory epithelial cell types becomes progressively restricted with age. (A) Experimental timeline showing tamoxifen induction and chase intervals. **(B)** Representative images of tdT^+^ progeny of *Ascl1*^+^ cells labeled at P0 and analyzed at weaning, including BG cells (left), Sus cells (middle), HBCs and neurons (right). Dotted lines indicate the basal lamina. Scale bar, 50 μm. **(C)** Representative images of lineage tracing from E14.5, P0, P3, P5, P8, or P35 showing an overall decline in labeled output with age. Scale bar, 200 μm. **(D)** Immunostaining for ASCL1 and PCNA at P1, P3, and P8. ASCL1^+^ cells appear more abundant and more likely to be PCNA^+^ at P3 than at P1 or P8. Dotted lines indicate the basal lamina. Scale bar, 50 μm. **(E)** Proportion of each lineage traced non-neuronal cell type among all non-neuronal cells at each developmental stage. Number at top of each bar indicates total cells counted. **(F)** Model summarizing progressive restriction of *Ascl1*^+^ cell fate capacity over time.

From embryonic and neonatal timepoints, *Ascl1^+^* cells generated apical SOX2⁺ sustentacular cells (Sus), basal SOX2⁺; P63⁺ horizontal basal cells (HBCs), and submucosal Bowman’s glands (BG) (**Fig. 1B**). Rare SOX9⁺ tdT⁺ microvillar (MV) cells were also observed (**Fig. S1**). Lineage-traced neurons, identified based on morphology and position in the OE, were more abundant than any other cell type at all ages (**Fig. 1C**). Neuron production dropped precipitously between E14.5 and P8, though there was a transient burst in neurogenesis around P3–P5 (**Fig. 1C**). To test the hypothesis that this elevated neurogenesis reflects increased proliferation of transit-amplifying neuronal precursors, we examined Proliferating Cell Nuclear Antigen (PCNA) abundance at P1, P3, and P8 and observed more PCNA^+^; ASCL1^+^ cells at P3 relative to P1 and P8 (**Fig. 1D**).

To assess the cell fate capacity of *Ascl1⁺* cells across development, we characterized tdTomato⁺ (tdT) cells using cell type-cell type-specific markers and analyzed the relative abundances of lineage-traced non-neuronal cells at different stages of development. We grouped the ages tested into four bins -- *Embryonic* (E14.5), *Early* (P0), *Middle* (P3, P5), and *Late* (P8, P35) -- and quantified the relative proportion of each non-neuronal fate in each bin (**Fig. 1E; Table S2**). Each BG/duct was counted as a single unit because it clonally derives from a cKit⁺ progenitor (Goss et al. 2016). We found that the relative proportions of HBCs, Sus cells, and BG/duct structures changed significantly over developmental time (Kruskal–Wallis, all p < 0.001): *Embryonic* progenitors generated the highest proportion of HBCs (*p* ≤ 0.024; BH-adjusted Wilcoxon pairwise test), *Late* progenitors generated the highest proportion of Sus cells (*p* ≤ 0.017), and relative BG/duct output was highest in the *Middle* stage (*p* ≤ 0.030). Interestingly, relative HBC production declined over time, while relative Sus production increased, though overall non-neuronal cell production was minimal by P8. Overall, these results suggest four major stages of *Ascl1^+^*non-neuronal cell fate capacity (**Fig. 1F**): **1)** a bias toward HBC fate during embryogenesis, **2)** a drop in HBC production by birth, **3)** a relative rise in Sus production in the first postnatal week, and **4)** another drop, to almost zero (**Table S2**), in overall non-neuron production by P8.

### Neuronal composition of the olfactory epithelial niche steadily increases over time

We next sought to identify extracellular and intracellular pathways that could affect these shifts in the fate capacity of *Ascl1*^+^ cells throughout development.

We performed single-cell RNA sequencing (scRNA-seq) of live cells isolated from the mouse OE on P1, P3, P5, P8, P15, and P22 (see Methods) (**Fig. S2A**). Removal of low quality cells from an initial 94,373 cells retained 82,677 cells (**Fig. S2B**). We performed unsupervised clustering using *Seurat* (Hao et al. 2024) and visualized the data using uniform manifold approximation and projection (UMAP) (**Fig. S2C-E**) (Blondel et al. 2008; Becht et al. 2018). The resulting 21 clusters were annotated using known marker genes (Brann et al. 2020; Gadye et al. 2017) (**Fig. S2F**, **Table S1**) and corresponded to sensory OE, non-sensory respiratory epithelium, stroma, immune cells, olfactory ensheathing cells, odontogenic epithelium, and olfactory bulb neurons (**Fig. S2F**). Cells from all ages and all sequencing batches were represented in all clusters except for clusters 5, 18, and 19, which lacked cells from one or more ages, and clusters 10, 17, and 20, which lacked cells from one or more mice (**Fig. S2C,D**).

Our lineage tracing results showed that the ratio of cell types produced by *Ascl1*^+^ cells shifts over time. We were therefore curious if the proportions of all cell types in the sensory OE changed throughout postnatal development. We first removed all cells not in the sensory OE lineage (**Fig. S3A,B**), then reclustered, visualized the results by UMAP (**Fig. 2A**), and annotated clusters using known marker genes (**Fig. 2C**) (see Methods). We then analyzed cluster composition by age and found that mOSNs progressively increased in relative abundance from P1 to P15 (14.3% at P1 vs. 53.1% at P22), while iOSNs and INPs were more abundant in younger animals, together accounting for 69.0% of cells at P1, but dropping on subsequent days (**Fig. 2B**). The proportion of GBCs peaked at 11.7% at P3 and then declined to 1.7% by P22.

**Figure 2:**
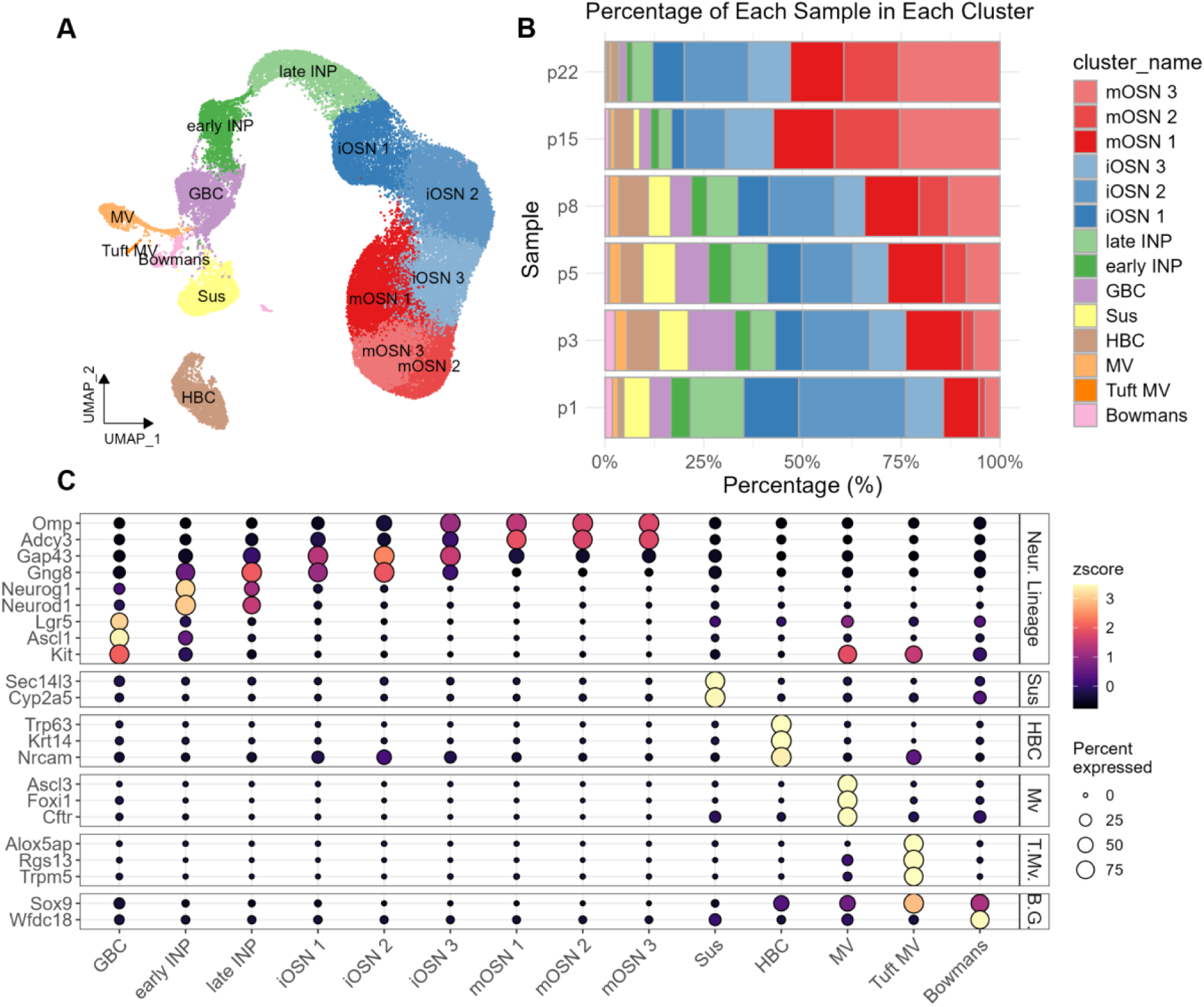
The composition of the sensory OE shifts from high to low relative abundance of non-neuronal cell types over postnatal development. **(A)** UMAP of sensory OE cells labeled by annotated cluster. **(B)** Stacked bar plot showing the percentage of cells at each age (P1, P3, P5, P8, P15, and P22) assigned to each cluster. **(C)** Dot plot of selected marker genes used for cluster annotation. Dot size indicates the fraction of cells expressing each gene, and color indicates scaled average expression.

Among non-neuronal lineage cells, Sus cells represented 58.4% and HBCs represented 12.4% at P1. By P22 however, their relative abundances switched, with Sus cells comprising only 11.9% of non-neuronal cells and HBCs comprising 43.3%. Tuft MV cells increased from birth to weaning (0.1% at P1 vs. 14.3% at P22), while BG cells decreased from 16.3% at P1 to 12.0% by P22. These ratios were largely consistent across replicates (**Fig. S3C; Table S1**).

### Subsets of *Ascl1*^+^ progenitor cells express canonical lineage-specific genes

To test the hypothesis that gene expression differences between individual *Ascl1^+^* cells might indicate commitment to specific lineages, we clustered the 4,478 cells expressing *Ascl1* (*Ascl1* > 0) using *Seurat* and visualized them by UMAP (**Fig. S4; Table S3**). We initially noticed a large group of cells that had relatively low expression of *Ascl1* but expressed canonical neuronal genes, such as *Ebf2*, possibly indicating cells committed to the neuronal lineage and down-regulating *Ascl1* expression; however, we were unable to confirm co-expression of ASCL1 and EBF2 protein using immunohistochemistry IHC (**Fig. S4**). The GBC-to-neuron lineage has been well characterized (reviewed in (Gokoffski et al. 2010)); therefore, for ease of visualization, and to focus on the non-neuronal lineages, we removed the five neuronal lineage clusters from further analysis.

We then used *Seurat* module scores for Sus, BG, HBC, MV, and GBC identity to characterize the seven remaining clusters. Cells with high scores for a specific module were enriched within discrete clusters (**Fig. 3A,B**). Feature plots also showed enrichment of cells expressing lineage-specific genes in discrete clusters, suggesting that these clusters may represent early lineage bias of sub-populations of progenitor cells (**Fig. 3C**). We refer to these *Ascl1*^+^ cells with high Sus, BG, MV, or HBC module scores as “transitional” cells, reflecting the co-expression of *Ascl1* with a marker of a non-neuronal lineage. Transitional cells were present at every time point, although they appeared to decrease in abundance after P8 (**Fig. 3D**). To verify that these are bona fide transitional cells and not artefacts (e.g., cell doublets during cell capture), we used IHC to validate their presence in vivo. We identified cells that co-express ASCL1 and P63 (putative HBC lineage), and ASCL1 and IL33 protein (putative Sus lineage) at P1 (**Fig. 3E,F**), and we observed occasional ASCL1; P63-double positive cells as late as P15 (data not shown).

**Figure 3:**
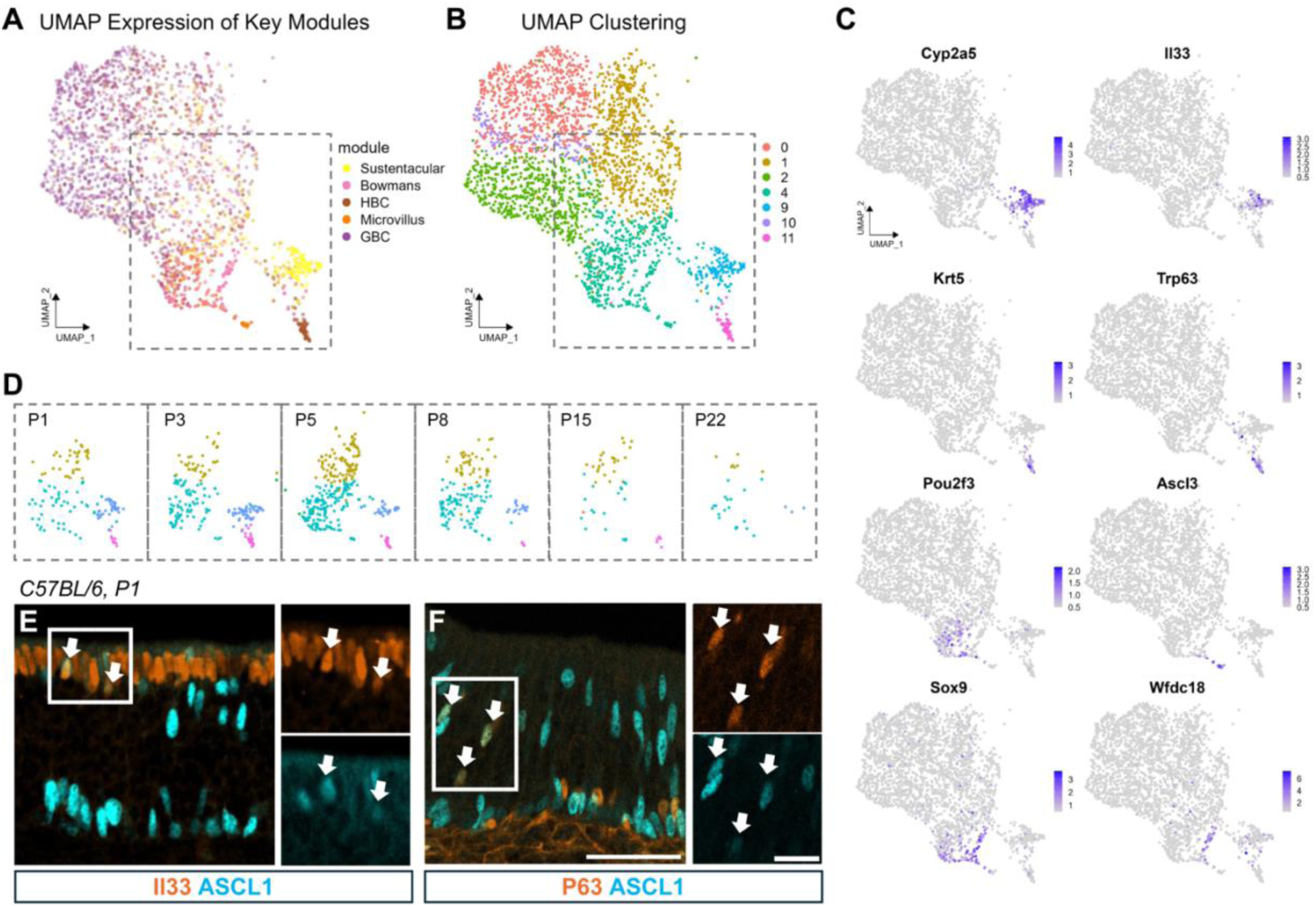
Clustering of *Ascl1*^+^ cells reveals early bias toward non-neuronal fates. **(A,B)** UMAP of *Ascl1*^+^ cells showing enrichment of cells scoring high for Sus, HBC, MV, and BG associated modules (A) in specific clusters (B). Module scores in (A) are plotted by overlaying cells with positive scores for each tested module on the shared UMAP with point color indicating module identity and point opacity scaled to module score. The dashed box marks the region enlarged in D; cells outside this region were omitted from the inset views for clarity. **(C)** Feature plots showing the expression of marker genes associated with distinct non-neuronal cell types. **(D)** Enlargement of the boxed region in A showing cells by age. Colors indicate cluster identity as in B. **(E,F)** Representative immunohistochemistry from animals at P1 confirms ASCL1 co-labeling with IL33 (E) and P63 (F). Scale bar 50 µm; Inset scale bar 20 µm.

### Trajectory analysis identifies non-neuronal lineages originating from *Ascl1*^+^ cells

Together, the above results suggest that *Ascl1^+^* cells generate all sensory OE cell types throughout the postnatal period. To identify pathways with a potential role in fate determination, we used the scRNA-seq data to reconstruct lineages arising from *Ascl1*^+^ cells. We limited our analysis to cells of the sensory OE but removed the BG and Tuft MV clusters due to low cell counts. Additionally, a subset of Sus cells showed high expression of neuronal genes and was excluded from the data set (see Methods) (**Fig. S5**, **Table S4**). To infer the developmental trajectory of *Ascl1^+^*cell progeny, we ran *slingshot* (Street et al. 2018) on PCA-reduced data. *Slingshot* identified a trajectory consisting of four lineages starting from *Ascl1*^+^ cells: HBC, Sus, MV, and mOSN (**Fig. 4A**).

**Figure 4:**
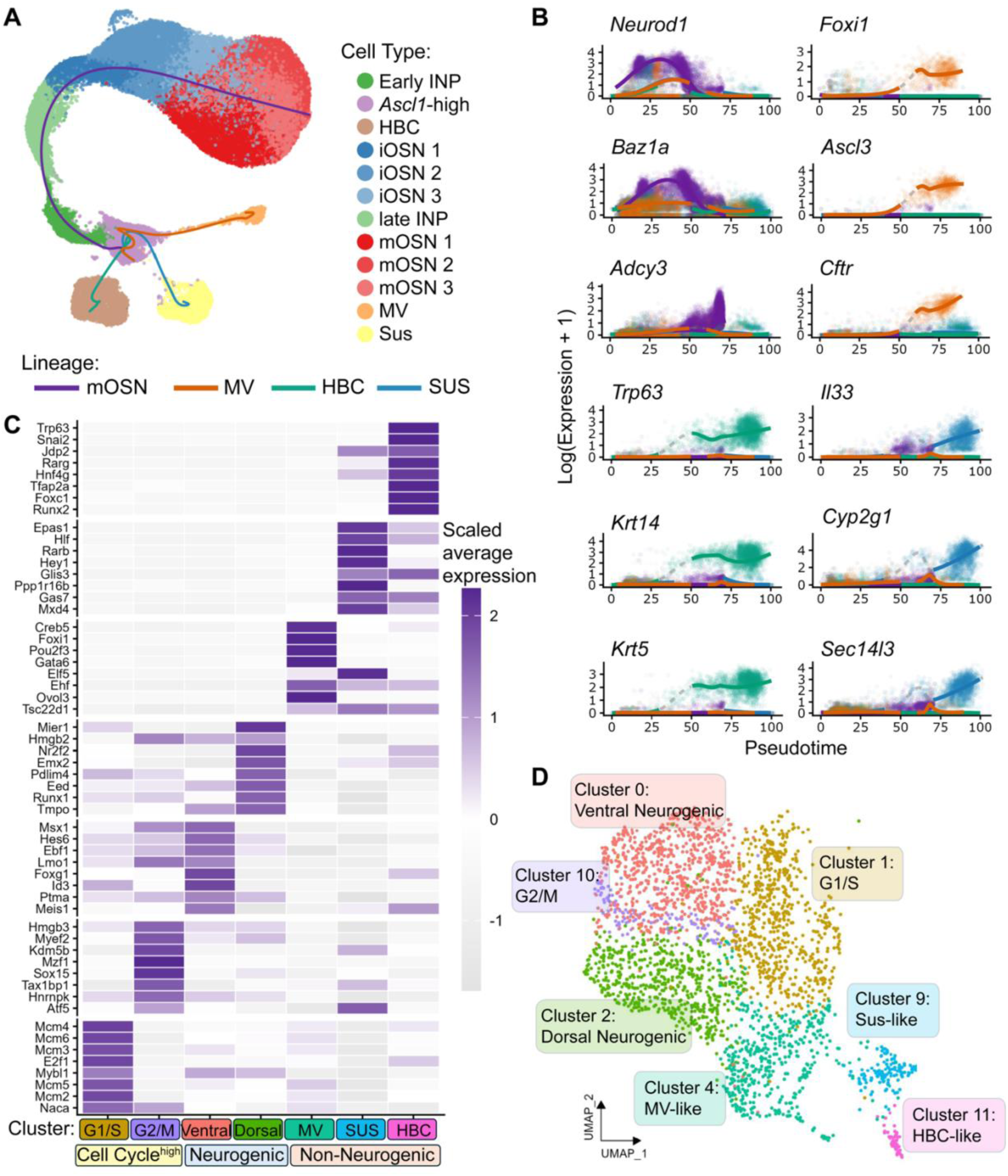
Differential expression along lineages arising from the *Ascl1*^+^ cells. **(A)** UMAP of sensory OE cells with the *Slingshot* trajectory overlaid showing four lineages starting in *Ascl1*^+^ cells and ending in HBCs, Sus cells, MV cells, or mOSNs. Cells are colored by annotated cell type, and curves defining the four lineage paths are colored by lineage. **(B)** Lineage-resolved gene expression across pseudotime. Points represent single cells and curves represent expression averaged over cells in each lineage. Dashed gray segments mark pseudotime intervals with fewer than 100 cells per 10 pseudotime units, where sparsity reduces confidence in the fitted trend. Neuronal lineage = purple, HBC lineage = brown, MV lineage = orange, Sus lineage = blue. **(C)** Heatmap showing the top 8 TFs for each cluster ranked by adjusted p-value and colored by scaled average expression. **(D)** UMAP of *Ascl1*^+^ cells colored by cluster and labeled by cluster annotation.

To better understand lineage differentiation, we used *tradeSeq* (Van den Berge et al. 2020), which identifies genes that change in expression as a function of pseudotime. We identified genes that increased significantly from the start to the end of a lineage and were at least 1 log-unit higher in a given lineage than in all other lineages at the end of the lineage (**Fig. 4B, Table S4**). Top genes upregulated along each lineage included canonical marker genes: *Trp63*, *Krt5*, *Krt14*, *Perp*, and *Nrcam* in the HBC lineage, *Cyp2g1*, *Ermn*, *Cyp2a5*, *Il33*, and *Sec14l3* in the Sus lineage, *Ascl3*, *Foxi1*, *Cftr*, and *Coch* in the MV lineage, and *Uncx*, *Ebf1*, *Gng8*, *Myt1l*, *Omp* and *Adcy3* in the mOSN lineage.

### Early transcriptional heterogeneity in *Ascl1*+ cells reveals candidate regulators of lineage divergence

We next returned to the *Ascl1*^+^ subclusters to ask whether transcriptional differences within this population might reveal candidate regulators of lineage divergence. We restricted our analysis to TFs, reasoning that TF enrichment may reflect early lineage-associated transcriptional programs (**Table S3**).

Using *Seurat* cluster markers, we found that the most enriched TF for the HBC-like cluster was *Trp63*, a key regulator of HBC quiescence (Fletcher et al. 2011; Packard et al. 2011) (**Fig. 4C, D**). Among other top TFs enriched in the HBC-like cluster were *Snai2*, *Foxc1*, and *Hnf4g*. Similarly, the MV-like cluster was enriched for *Pou2f3*, which is required for specification of MV and BG cells (Yamaguchi et al. 2014; Weng et al. 2016), as well as *Gata6* and *Ehf*. Enriched TFs in the Sus-like cluster included the Notch effector *Hey1*, the Sus marker *Epas1* (Nickell et al. 2012), and *Glis3*. Non-differentiated-like clusters resembled neurogenic and/or cycling progenitor states: clusters 0 and 2 showed ventral and dorsal neurogenic signatures, respectively (*Foxg1*, *Acsm4*), while cluster 1 was enriched for transcripts associated with G1/S phases (*Mcm5*, *Cdc6*), and cluster 10 showed high expression of transcripts associated with G2/M phases (*Cdc20*, *Ccnb2*) and self-renewal/repression of alternative lineages (*Kdm5b*) (Xie et al. 2011; Schmitz et al. 2011) (**Fig. 4C, D**).

These data suggest that lineage divergence of *Ascl1*^+^ cells may occur early, while cells are still cycling and expressing *Ascl1*, and may be driven by known and novel fate specification transcriptional programs.

### *Trp63* plays a role in HBC lineage commitment

To address the hypothesis that transitional cells represent progenitors biased toward specific fates, we tested the role of the gene encoding tumor protein P63 (*Trp63*) in HBC fate specification. Previous studies have shown that mice lacking *Trp63* from germline do not produce HBCs (Packard et al. 2011), and specific ablation of *Trp63* from adult HBCs releases them from quiescence, although they lose the capacity for self-renewal (Fletcher et al. 2011, 2017). Our *tradeSeq* analysis showed that *Trp63* was strongly associated with the HBC lineage and increased consistently from the start to the end of the lineage (**Fig. 4C, Table S4**). In addition, *Trp63* was the most significantly enriched TF in the HBC-like cluster, suggesting it may play a role in the transition from *Ascl1*^+^ progenitor to mature HBC. To specifically reduce *Trp63* in *Ascl1^+^* cells during OE development, we used timed-pregnant *Ascl1^CreER^; Trp63^lox/+^; R26^tdT^* (“*Trp63*-het”) mice and administered a single dose of tamoxifen to induce Cre-mediated recombination at E14.5, around the time that *Ascl1^+^* cells first start making HBCs (Packard et al. 2011). When we collected the OE from embryos at E19.5, we found that a smaller fraction of HBCs were lineage-traced in *Trp63*-het animals compared with wild-type controls (*p*-value = 0.031), suggesting that some *Ascl1*^+^ cells that would normally give rise to HBCs were diverted to alternative fates (**Fig. 5; Table S2**).

**Figure 5:**
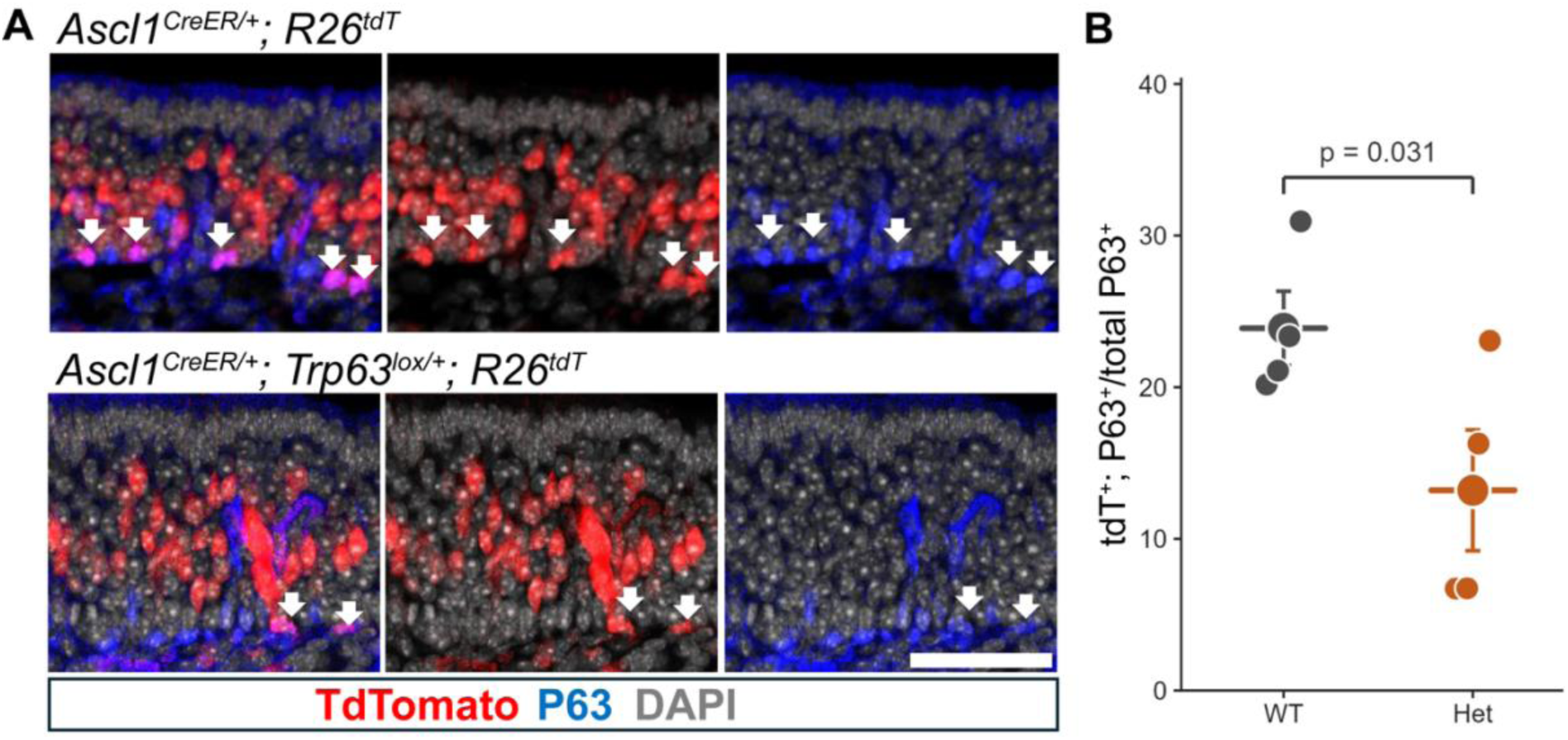
*Trp63* knockdown in *Ascl1*^+^ cells reduces the percentage of lineage-traced HBCs among all HBCs. **(A)** Lineage-traced cells in OE of *wild-type* (top) or *Trp63-heterozygous* (bottom) embryos induced at E14.5 and dissected at E19.5. Scale bar, 50 μm. **(B)** Percentage of HBCs that were lineage-traced in *wild-type* versus *heterozygous* animals. N = 4 mice per group.

### Perinatal HBCs are proliferative but limited to self-renewal

Adult HBCs are multipotent reserve stem cells that can regenerate all OE cell types after acute injury (Iwai et al. 2008; Leung et al. 2007). Their contribution to normal OE development, however, is still unclear. To identify the cell fate capacity of perinatal HBCs, we crossed mice expressing a tamoxifen-inducible Cre transgene driven by the bovine *Keratin 5* promoter (Indra et al. 1999), which in the OE is specific to HBCs, with *R26^tdT^* mice. A single dose of tamoxifen was administered to *Krt5-CreER(T2)*; *R26^tdT^* pups on P1, P3, P5, or P8 -- all of which were collected at weaning -- or to young adults at P33-P34, which were collected at P47-P48 (**Fig. 6A**). In contrast to *Ascl1^+^*progenitor cells, *Krt5^+^* progenitor cells were restricted to the HBC lineage; despite robust expression of the S-phase marker PCNA in neonatal HBCs (**Fig. 6B**) and large numbers of HBCs identified as being in S-or G2/M phase **(Fig. 6C)**, their progeny overwhelmingly consisted of other HBCs (**Fig. 6D**). Extremely rare patches of mOSNs and Sus cells were observed at all timepoints, and a few tdT^+^ BG were present after P8 induction in one mouse; however, occasional differentiation is consistent with the known role of HBCs in homeostatic turnover (Fletcher et al. 2011, 2017).

**Figure 6:**
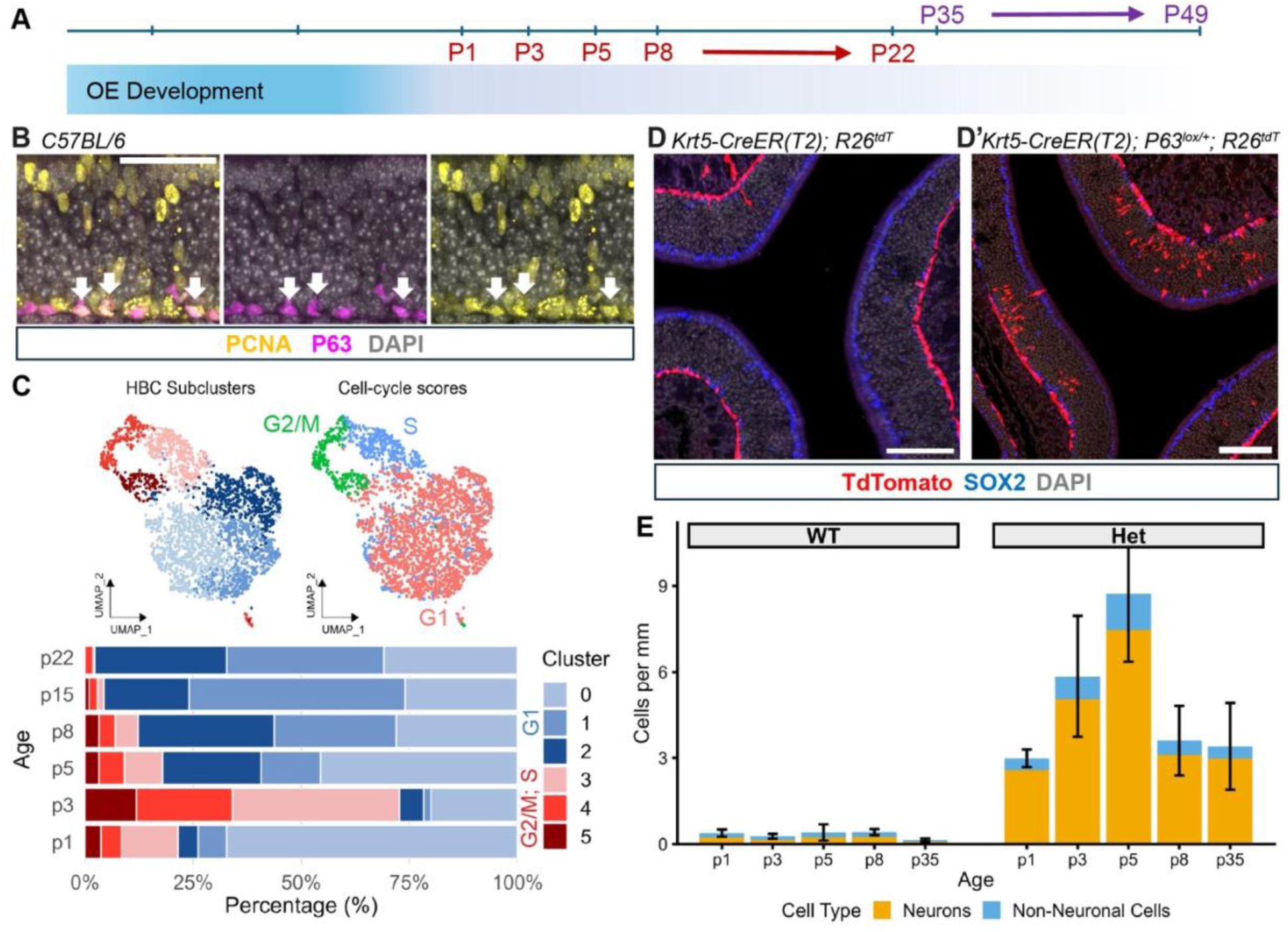
*Trp63* restricts HBC fate in a dosage-dependent manner. **(A)** Timeline showing tamoxifen induction and chase intervals. **(B)** A subset of HBCs, identified by P63 (magenta) and laminar position, express PCNA (yellow) at P0. DAPI shown in grey. Scale bar = 50µm. **(C)** UMAPs colored by cluster (left) or cell-cycle phase assignment (right). HBCs in S and G2M phases are enriched at early postnatal timepoints and decline with age. Stacked bar chart (bottom) shows relative proportion of cells in each cluster at each time point. **(D,D’)** Representative images of tdT^+^ progeny generated from *Krt5-CreER(T2); R26^tdT^* (D) animals and *Krt5-CreER(T2); p63^lox/+^;* R26^tdT^ (D’) animals induced at P5. Scale bars = 100 μm. **(E)** Density (cells per mm of OE) of tdT^+^ neurons (yellow) and non-neuronal cells (blue) produced by *Trp63^+/+^* (left) and *Trp63^fl/+^* (right) HBCs at each age of induction. Error bars indicate SEM.

### Loss of one copy of *Trp63* expands perinatal HBC lineage capacity

The loss of one copy of *Trp63* in adult HBCs leads to occasional differentiation of HBCs into Sus cells or mOSNs (Fletcher et al. 2011; Fletcher et al. 2017). To address the hypothesis that neonatal HBCs, while fate-restricted to self-renewal, are sensitive to P63 levels, we deleted one copy of *Trp63* in developing and adult HBCs and compared the identity of lineage traced cells with that of cells traced from WT HBCs. P63 knockdown was induced in *Krt5-CreER(T2)*; *p63^lox/+^*; *R26^tdT^*animals using a single dose of tamoxifen at P1, P3, P5, P8, or P35. Pups were dissected for analysis at weaning, and adults induced at P35 were dissected at P49. Images of the OE were then scored for the number of lineage-traced differentiated cells (non-HBCs) (**Fig. 6D’**).

We observed markedly more lineage-traced differentiated cells in *Trp63*-het animals compared with WT controls (**Fig. 6E**). To assess the effect of genotype and age on HBC differentiation, we fit separate additive linear models with genotype and age as predictors. For neuronal differentiation, genotype showed a strong effect (F(1,27) = 96.68, *p* = 2.04×10^-10^), whereas age did not (F(4,27) = 1.44, *p* = 0.248). For non-neuronal differentiation, genotype again showed a strong effect (F(1,27) = 29.19, *p* = 1.04×10^-5^), while age did not (F(4,27) = 2.23, *p* = 0.092). Adding a genotype-by-age interaction did not significantly improve model fit for either neuronal density or non-neuronal density. Together, these results indicate that *Trp63* heterozygosity expanded the capacity of HBCs to generate neuronal and non-neuronal cells, independent of age. We note that in comparison with previous results (Fletcher et al. 2017) we observed more frequent differentiation of *Trp63*-het HBCs, which may be the result of differences in aspects of the model used, such as fluorescent marker (YFP vs. tdT).

### Cell-cell signaling within the stem cell niche evolves over developmental time

Shifts in the cell fate capacity of *Ascl1^+^* cells correspond with changes in the numbers and types of cells present in the OE niche. We therefore wanted to identify potential extracellular signaling pathways that could feed back onto *Ascl1^+^* cells to affect cell fate and to better understand how these stem cells may integrate extracellular signals with intracellular regulation of gene expression.

We used *CellChat* (version 2.2.0) (Jin et al. 2021) to computationally identify possible ligand-receptor pairs between GBCs and other cell types in the niche (**Fig. 7; Table S5**). Signaling to GBCs appeared highest at P3, remained elevated at P5, and appeared weakest at P22, dropping in predicted strength by 89% between P5 and P22 (**Fig. 7A**). Among the 12 sender cell types we analyzed (mOSN, iOSN, INP, Sus, HBC, GBC, fibroblasts, respiratory basal cells, MV, respiratory goblet cells, inflammatory monocyte-like cells, and macrophage-like cells) (**Fig. S2F**), cells of the sensory OE (mOSNs, Sus, INPs, iOSNs, HBCs, and GBCs) contributed the strongest signals to GBCs at all stages of development (**Fig. S6**). Comparison of individual pathway strength between P5 and P22 predicted a reduction in MK, PTN, ADGRL, and CADM signaling to GBCs from multiple sender populations (**Fig. 7B**). Some of these pathways share common receptors, such as Nucleolin (*Ncl*) and Syndecan-4 (*Sdc4*). Top ligands predicted for HBC-to-GBC signaling, specifically, included *Ptn*, *Mdk*, *Tenm3/4*, and *Kitl* (**Table S5**). Overall, these changes indicate a shift from a highly proliferative, highly trophic early niche dominated by growth factor and adhesion signaling to a more restricted, less plastic niche in adulthood.

**Figure 7.**
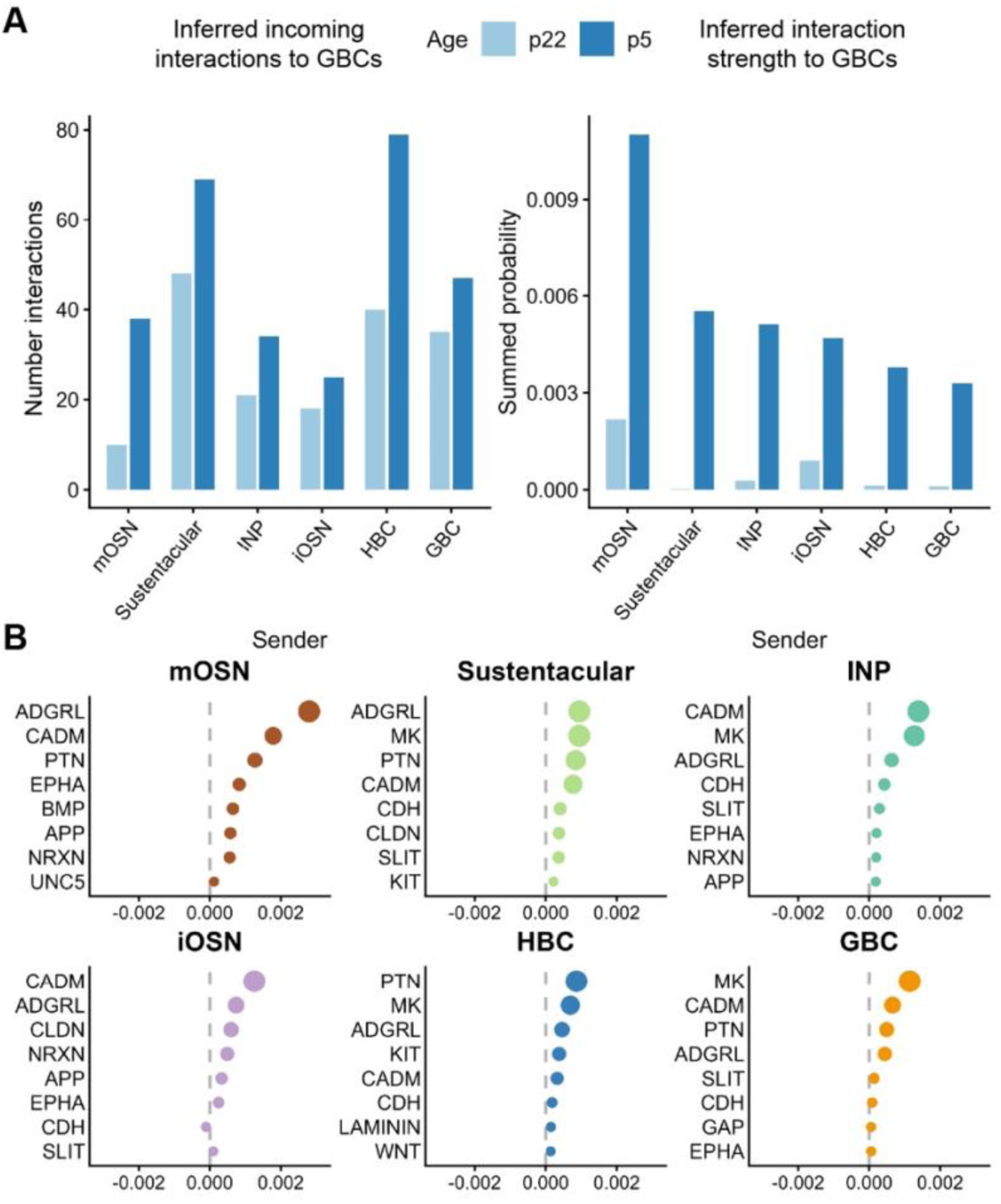
Developmental changes in niche signaling to GBCs. **(A)** Bar graphs showing incoming signaling to GBCs, represented by the number of inferred incoming interactions (left) or the summed *CellChat* interaction strength (right) for the top six sender cell classes at P5 (dark blue) and P22 (light blue). Additional analyzed sender classes with weaker predicted input are shown in Supplementary **Fig. S6**. **(B)** Most changed signaling pathways for each sender to GBCs, plotted as pathway-level differences in *CellChat* probability between P5 and P22. Points to the right of zero indicate pathways stronger at P5; points to the left of zero indicate pathways stronger at P22. Point size reflects the magnitude of the difference.

## Discussion

In the present study, using genetic lineage tracing, we provide direct evidence that the lineage capacity of *Ascl1*^+^ cells in the OE changes over developmental time, from relatively high production of HBCs during the embryonic period to almost exclusive production of neurons by adulthood. By contrast, *Krt5*^+^ HBCs are lineage-restricted throughout postnatal development despite being proliferative and multipotent. Single-cell RNA sequencing enabled us to reconstruct lineages arising from *Ascl1*^+^ cells, revealing transitional cell states between multipotent and committed progenitors. We also identified potential cell-cell communication pathways that may signal from daughter cells back to progenitor cells.

### Evolving niche factors may influence the developmental capacity of *Ascl1*^+^ cells

In other regions of the nervous system, progenitor cells stereotypically generate different cell types at different times, indicating developmental shifts in cell fate capacity. Studies have shown that cell-extrinsic niche signaling shapes fate decisions in stem cells by integrating with cell-intrinsic transcriptional changes to ensure the proper balance of cell types as the niche develops (Beumer and Clevers 2024; Enver et al. 2009; Rompolas et al. 2013). During cortical development for example, cortical progenitors become progressively restricted in their competence to respond to external cues, generating deep layer neurons first and each successive upper layer at progressively later time points (McConnell 1995; McConnell and Kaznowski 1991; Frantz and McConnell 1996). Similarly, retinal progenitor cells produce the seven major retinal cell types in a sequential and overlapping manner through a combination of cell-intrinsic and -extrinsic mechanisms (Yang 2004; Morrow et al. 1998; Belliveau and Cepko 1999).

Although it is still unclear whether and to what extent the developmental potential of OE progenitor cells is determined intrinsically and to what extent the local microenvironment affects cell fate, our data show that the relative proportions of cell OE types -- and therefore potential inputs from these cells to progenitors -- shifts over time. Extracellular signaling to GBCs was predicted to be higher at P5 than at P22, suggesting a highly trophic environment in young OE, with high Midkine (MK), pleiotropin (PTN), and ADGRL signals from Sus cells, GBCs, and HBCs. MK and PTN are heparin-binding growth factors expressed during morphogenesis of a variety of tissues (Muramatsu 2002), and ADGRL is an adhesion G-protein coupled receptor linked to tissue development, growth, and organization (Sreepada et al. 2022). A recent study reported that HBCs provide a supportive niche for GBC survival and neurogenesis in part via MK-SDC4, KITL-KIT, and WNT signaling (Gameiro et al. 2025). Consistent with this, our *CellChat* analysis predicted HBC-to-GBC signaling via *Mdk*-*Sdc4*, *Kitl*-*Kit*, and *Wnt4/6*-*Fzd2/3/8*-*Lrp5/6* interactions.

### Cell-intrinsic mechanisms contribute to lineage restriction

Cell-intrinsic mechanisms, such as epigenetic modulators or transcription factors, may interact with niche signals to affect the cell fate capacity of progenitor cells. The histone methyltransferase EZH2, for instance, restricts basal fate in lung and muscle development (Snitow et al. 2015, 2016), and the transcription factor FOXG1 represses Cajal-Retzius (C-R) fate during cortical development; its depletion allows cortical progenitors to continue making C-R cells beyond the normal timeframe for C-R production (Hanashima et al. 2004). It is still unclear, however, how these internal programs ensure fate restriction. We show, for instance, that HBCs continue to cycle well after committing to HBC fate but are restricted to the production of HBCs, a characteristic that persists throughout the changing niche. Interestingly, reduction of *Trp63* in HBCs at all ages tested expanded their capacity to produce alternative cell types, suggesting that *Trp63* may act in part to repress lineages otherwise permitted by the microenvironment. In contrast to HBCs, *Ascl1^+^* cells become gradually but progressively restricted to the neuronal lineage over time and retain some ability to make non-neuronal cells after injury, suggesting that crosstalk between cell-intrinsic factors and the microenvironment may play a role in lineage capacity (Goldstein et al. 1998; Peterson et al. 2019; Lin et al. 2017; Huard et al. 1998; Leung et al. 2007; Schnittke et al. 2015).

### Transitional cells embody a primed stem cell state

Progenitor cells can co-express TFs associated with distinct lineages or states, a phenomenon that has been variously described in the literature as lineage plasticity, promiscuity, or priming (Redó-Riveiro et al. 2024; Wuenschell et al. 1996; Hu et al. 1997). In early development of the mouse lung, for instance, distal airway epithelial progenitor cells co-express markers of neuroendocrine, clara, and type II alveolar cells before later differentiating into these separate lineages and resolving their gene expression to match (Wuenschell et al. 1996). Similarly, hematopoietic progenitor cells co-express TFs characteristic of multiple lineages, and gradual shifts in the balance of these TFs reversibly drives cell fate (Palii et al. 2019).

In the present study, we refer to *Ascl1*^+^ cells that co-express markers of non-neuronal lineages as “transitional” to account for the possibility that ASCL1 may act as a differentiation signal rather than a neuronal determination signal. Consistent with this interpretation, we found expression of TFs in subclusters of *Ascl1*^+^ cells that may reflect transcriptional priming of these cells toward specific non-neuronal lineages. *Elf5* and *Glis3*, for instance, which have been shown to specify cell fate in breast, pancreas, or sperm (Oakes et al. 2008; Lee et al. 2011; Chakrabarti et al. 2012; Li et al. 2019; Scoville et al. 2017), were expressed in the Sus-like subcluster. Similarly, the MV-like subcluster was enriched for *Gata6*, *Ehf*, and *Klf4*, which are required for cell fate specification and/or regeneration in lung or cornea (Yang et al. 2002; Keijzer et al. 2001); (Stephens et al. 2013). In the HBC-like subcluster, we identified *Snai2*, a classic marker of epithelial-mesenchymal transition (Mistry et al. 2014; Wei et al. 2020) the pioneering TF *Foxc1* (Xu et al. 2021; Ray et al. 2010; Wang et al. 2016), and *Hnf4g*, a key regulator of intestinal maturation (Chen et al. 2021; Li et al. 2019).

Transcriptional priming in OE progenitor cells may follow epigenetic priming of lineage-specific genes, which could allow for quick response to a differentiation cue. In embryonic stem cells (ESCs), the promoters of differentiation genes have both active and repressive histone modifications (so-called “bivalent domains”), often form complexes with poised RNA polymerase II, and may exhibit co-occupancy of the transcriptional repressor SUZ12 and an ESC transcriptional activator (OCT4, SOX2, and/or NANOG), suggesting that they exist in a primed pre-differentiation state (Azuara et al. 2006; Bernstein et al. 2006; Lee et al. 2006; Stock et al. 2007). Similar stem cell priming has been observed in adult stem cell populations (Reizel et al. 2021; Norrie et al. 2025), including adult HBCs (Van den Berge et al. 2026). In developing human cerebral cortex, chromatin accessibility around lineage specific genes, particularly TFs, precedes gene expression in subsets of cycling progenitors, suggesting that they are epigenetically primed for specific lineages (Trevino et al. 2021). Interestingly, one such lineage consists of an early multipotent glial progenitor state with the capacity to differentiate into both astrocytes and oligodendrocytes and co-expresses ASCL1, EGFR (astrocyte lineage) and OLIG2 (oligodendrocyte lineage) protein, consistent with transcriptional priming of multiple lineages.

Primed progenitor cell states may allow for lineage plasticity, or the ability of a cell to be diverted to an alternative fate. For example, in dopaminergic neurons differentiated from ESCs, non-neuronal lineage TFs remained bound by RNA polymerase II, perhaps allowing for potential reactivation (Ferrai et al. 2017). In the perinatal olfactory epithelium, *Trp63* was the most enriched TF in the HBC-like *Ascl1*^+^ subcluster; co-expression of P63 and ASCL1 in a subset of cells suggested that P63 may prime *Ascl1*^+^ progenitors for the HBC developmental program. Consistent with this view, deletion of one copy of *Trp63* in *Ascl1*^+^ cells at E14.5 appeared to shift some cells away from HBC fate. Together, these data illustrate transcriptional priming and lineage plasticity of *Ascl1*^+^ cells during a highly trophic developmental window between the embryonic period and first postnatal week, after which these cells become restricted to a single lineage under homeostatic conditions.

## Acknowledgments

The authors would like to thank members of the Ngai, Purdom, and Dudoit labs, especially Shelby Jones, Amy Cao, Jonathan Lovas, Alvin Tak, and Brittany Brooks for ongoing discussions and technical support. This work used the computational resources of the National Institutes of Health (NIH) HPC Biowulf cluster (https://hpc.nih.gov) and the National Heart, Lung, and Blood Institute DNA Sequencing and Genomics core facility and was supported by the Division of Intramural Research of the National Institute of Neurological Disorders and Stroke, NIH (ZIA NS009423) (J.N.). The content is solely the responsibility of the authors and does not necessarily represent the official views of the NIH.

## Materials and Methods

### Animals

All animal work was carried out in compliance with the National Institute of Neurological Disorders and Stroke ACUC (ASP 1579) according to federal guidelines. *Ascl1*-*CreER* knock-in mice (Kim et al. 2011), *R26^tdTomato^*reporter mice (Madisen et al. 2010), *R26^YFP^*reporter mice (Srinivas et al. 2001), *Krt5-CreER(T2)* transgenic mice (Indra et al. 1999), and mice harboring the *p63^lox^* allele (Mills et al. 2002) were bred and maintained on a C57Bl/6 background. C57Bl/6 mice were obtained from The Jackson Laboratory. Both male and female mice were used in all studies, although the P5 scRNA-seq sample consisted solely of males. The genotypes of mice used were:

*Ascl^CreER/+^; R26^Ai9/+^* (lineage tracing)

*Ascl^CreER/+^; R26^Ai9/+^*; *p63^lox/+^*(genetic manipulation of *Ascl1*^+^ cells)

*Krt5-CreER(T2)-Tg; R26^Ai14/+^* (lineage tracing)

*Krt5-CreER(T2)-Tg; R26^Ai14/+^*; *p63^lox/+^*(genetic manipulation of HBCs)

*Krt5-CreER(T2)-Tg; R26^YFP/YFP^*(single cell sequencing)

*R26^Ai14/+^*(single cell sequencing)

*Wild-type (C57Bl/6)* (single cell sequencing)

### Lineage tracing

#### Embryonic induction

To induce loxP recombination in embryos, timed-pregnant *Ascl^CreER/+^;R26^Ai9/+^* or *Ascl^CreER/+^;R26^Ai9/+^*;*p63^lox/+^* mice were given an intraperitoneal (IP) injection of tamoxifen (20 mg/mL dissolved in sterile corn oil) at a dose of 100 mg/kg on embryonic day (E)14.5. The morning of the vaginal plug was designated E0.5. On E19.5, the dam was euthanized and embryos were extracted for sample collection.

### Postnatal day 0 (P0), P1, P3, P5, and P8 induction

To induce loxP recombination in neonatal *Ascl^CreER/+^;R26^Ai9/+^, Krt5-CreER(T2)-Tg; R26^Ai14/+^*; *p63^lox/+^, or Krt5-CreER(T2)-Tg; R26^Ai14/+^* mice, 50 μg tamoxifen (Sigma Aldrich, T5648) dissolved in 50 μL sterile corn oil was injected intragastrically into the milk spot (Pitulescu et al. 2010; Lizen et al. 2015; Bakalar et al. 2023). *Adult induction*

To induce loxP recombination in adult *Ascl^CreER/+^;R26^Ai9/+^, Krt5-CreER(T2)-Tg; R26^Ai14/+^*; *p63^lox/+^, or Krt5-CreER(T2)-Tg; R26^Ai14/+^* mice, animals were given IP injections of tamoxifen dissolved in sterile corn oil at a concentration of 20 mg/mL at a dose of 100 mg/kg.

### Histology

Tissue was prepared for IHC as described in (Van den Berge et al. 2026) with the following modifications: EDTA-mediated decalcification was not used for animals P5 and younger. Frozen sections were sliced to 15 µm using a cryostat (Leica Biosystems), and slides were stored at −30°C or −80°C until use. When removed from the freezer, slides were dried on the bench, then washed in 1x PBS for 5 minutes on a shaker. Slides were again dried at room temperature for 30 minutes. For antigen retrieval, slides were immersed in Coplin jars filled with pre-heated 1x sodium citrate buffer pH 6.0 and placed in a steamer for 5 minutes. Coplin jars were then removed from the steamer and allowed to cool at room temperature for 30 minutes on the bench then washed in dH2O for 2 minutes. Next, slides were blocked in 1x PBS + 0.1% Tween-20 (PBST) containing 10% Normal Goat Serum (NGS), or Normal Donkey Serum (NDS) when Goat primary antibodies were used, for 1.5-2 hours. Slides were incubated in primary antibodies diluted in PBST containing 1% NGS/NDS at 4°C overnight (**Table S6**). The following day, the primaries were washed 3 x 5 minutes in 1X PBS on a shaker. Slides were incubated in secondary antibodies diluted in PBST + 1% NGS/NDS for 1-2 hours then washed 3 x 5 minutes in 1X PBS. Sections were mounted with glass coverslips using Fluoromount-G (Invitrogen, 00-4958-02) or ProLog Gold (Invitrogen, P36934) and sealed with clear nail polish. Sections were imaged at 20x magnification on a Stellaris 8 confocal microscope (Leica Microsystems).

### Image quantification and statistical analysis

#### Cell type identification

To quantify the progeny of *Ascl1*^+^ progenitor cells (**Fig. 1**) and differentiating *Trp63-*heterozygous cells (**Fig. 6**), tdT-labeled cells were classified into one of six categories based on antibody colocalization and cellular morphology: (1) HBC, (2) mature sustentacular cell, (3) Bowman’s gland, (4) microvillus cell, (5) neuron, and (6) immature sustentacular cell. Mature sustentacular cells were SOX2-positive and localized to the apical layer of the olfactory epithelium (OE). Immature sustentacular cells also expressed SOX2 but were positioned in the intermediate zone between the basal and apical layers. Bowman’s glands were SOX9-positive and identified in the submucosal layer. Each Bowman’s gland/duct was counted as a single unit because it represented a clonal structure derived from a cKit⁺ progenitor (Goss et al. 2016). Microvillus cells were SOX9-positive and located within the apical layer. HBCs were identified by basal positioning, morphology, and the expression of P63. Neurons were primarily found in the intermediate zone and were identified by morphology rather than by colocalization with antibody staining.

### Cell Counting

To quantify developmental changes in cell production from *Ascl1*^+^ progenitors and from *Trp63* heterozygous cells, we counted the total number of tdT+ non-neuronal cells in the sensory (non-respiratory) olfactory epithelium (OE) that were lineage traced from each developmental time point and normalized counts to the length of the olfactory epithelium basal layer in mm (cells per mm). OE length was measured using the *Fiji* (Schindelin et al. 2012) plug-in NeuronJ (Meijering et al. 2004). NeuronJ uses local image-intensity changes to identify likely epithelial boundaries, followed by user-guided live-wire tracing of the OE contour. For *Ascl^CreER/+^; R26^Ai9/+^* tracing we did not quantify neurons, although differences in production between timepoints had high “interocular significance” (Edwards et al. 1963). Cell counting was performed manually in *Fiji* using the Multi-Point tool, with a minimum of three images per animal averaged to produce a final score for that animal. From these data, cells per mm of OE and relative abundance of each cell type were calculated (**Table S2**).

### Analysis of Ascl1^+^ progeny over time

To assess developmental changes in the relative composition of non-neuronal progeny of *Ascl1*^+^ progenitors over time, we calculated, for each sample, the percentage of lineage-traced non-neuronal cells that were HBCs, Sus, or gland/duct cells (**Fig. 1C; Table S2**). Ages were binned into four developmental stages (Embryonic = E14.5; Early = P0; Middle = P3–P5; Late = P8–P35). For each cell type, we tested the null hypothesis that its relative proportion did not differ across developmental stages using a Kruskal-Wallis rank-sum test (kruskal.test). When the omnibus test was significant, pairwise differences between stages were assessed using

Wilcoxon rank-sum tests with Benjamini-Hochberg adjustment for multiple comparisons (pairwise.wilcox.test, p.adjust.method = “BH”, exact = FALSE).

### Analysis of the effect of Trp63-knockdown

To assess the effect of *Trp63* reduction on the fate of *Ascl1*^+^ cells (**Fig. 5; Table S2**), we counted all P63+ HBCs visible in the sensory OE of each embryo and then calculated the proportion of these HBCs which were lineage traced. Genotypes were compared using a one-tailed Student’s t-test. Note that the WT embryos used in the embryonic *Trp63* knockdown experiment are the same WT embryos contributing to the embryonic *Ascl1* tracing dataset.

To assess the effect of *Trp63* reduction on the fate of *Krt5*^+^ cells over time (**Fig. 6; Table S2**), we analyzed neuronal and non-neuronal cell densities using linear models. For each mouse, neuronal density was defined as lineage-traced neurons per mm of OE, and non-neuronal density was calculated as total lineage-traced cells per mm minus neuronal cells per mm of the OE. Because both values were skewed and contained zeros, each was transformed as log(1 + x) before modeling where log is natural logarithm.

For each outcome, we fit an additive ordinary least squares linear model of the form log(1 + response) ∼ Genotype + Age in R. Genotype (WT, Het) and age (P1, P3, P5, P8, P35) were modeled as categorical predictors, with WT as the reference genotype and P1 as the reference age. We also tested whether the effect of genotype varied by age by fitting models with a Genotype x Age interaction and comparing them to the additive models using nested-model F tests. The interaction term did not significantly improve model fit for either neuronal density (F(4,23) = 1.38, p = 0.271) or non-neuronal density (F(4,23) = 1.85, p = 0.153), we report the additive models. Overall effects of genotype and age were assessed from the additive models and coefficient estimates, standard errors, t-statistics, and p-values were extracted from the model summaries (Table S2).

### Cell capture and library preparation

The sensory OE was microdissected from the mouse nasal cavity at P1, P3, P5, P8, P15, and P22, dissociated in 2-3 mL pre-warmed dissociation medium (Worthington) (150 units papain, 100 units DNAse I, 2.5 mM cysteine, and 2.5 mM EDTA in 5 mL Neurobasal), and kept at 37°C for 25 min in a CO2 incubator. Samples were washed 3 times with 10% fetal bovine serum in phosphate-buffered saline (FBS-PBS), gently triturated 5-10 times using a fire-polished glass Pasteur pipette, and strained through a 35 μm nylon filter into a 5 mL polypropylene tube. DAPI, used to identify dead cells, was added at a final concentration of 0.1-1 μg/mL, and live cells (DAPI-low) were collected using a Sony LE-SH-800G cell sorter. We included at least two biological replicates (a biological replicate corresponds to an individual mouse or, rarely, a pair of sex-matched littermates) and at least one female and one male per time point for all time points except P5, which contained only males (**Fig. S2**).

Cells were pelleted in a swinging bucket centrifuge at 400g for 5 minutes at 4°C. The cell pellet was resuspended in 43.2 μl FBS-PBS, and 31.8 μl RT master mix was added to the cell pellet before loading the cells onto a Next GEM Chip G. A Chromium Next GEM Single Cell 3′ Kit v3.1 (10x Genomics, PN-1000268) and Dual Index Kit TT Set A (10x Genomics, PN-1000215) were used to generate single-cell whole-transcriptome libraries according to the manufacturer’s instructions.

### Sequencing

Indexed single-cell libraries were sequenced in three multiplexed batches on Illumina Novaseq 6000 or Novaseq X sequencers to produce paired-end reads. Fastq files were generated from binary base call files, aligned to mm10, and quantified using Cell Ranger v9.0.1 (Zheng et al. 2017). Output files consisted of a feature-barcode matrix containing a unique molecular identifier (UMI) for each transcript molecule associated with a feature/gene (row) and barcode/cell (column) and a molecule information file containing the number of reads assigned to a given feature for each cell.

### scRNA-seq data analysis

#### Preprocessing

Preprocessing and clustering were performed in *Seurat* v5 (Hao et al. 2024). Feature-barcode matrices produced by *Cell Ranger* were imported using Seurat::CreateSeuratObject with minimum features set to 200. Quality control was performed on the unintegrated layers of the Seurat object: after retaining cells with nUMI >= 1000, nGene >= 1000, log10GenesPerUMI > 0.8, and MitoRatio < 0.20, 82,677 cells remained from an initial 94,373 cells for downstream analysis (**Fig. S2**; **Table S1**). On average, we detected 3,643 genes and 11,767 unique molecular identifiers (UMIs) per retained cell. Subsequent analyses were performed as described below. Key software packages, version information, and computational resources are listed in **Table S6**.

### Normalization, PCA, clustering, and annotation

For all downstream analyses, subsets of cells were analyzed as follows: Data were log-normalized using Seurat::LogNormalize with a scale factor of 10,000, and 2,000 highly variable genes were identified using the vst method. Scaled expression values were used for principal component analysis (PCA). The number of principal components (PCs) used downstream was selected algorithmically as the minimum of two heuristics: the first PC at which cumulative variance exceeded 90% while individual PC variance fell below 5%, and the first PC after which the drop in variance between successive PCs was less than 0.1%. Using the selected number of PCs, nearest neighbors were identified with Seurat::FindNeighbors (k.param = 20; default Annoy search with Euclidean distance), and shared nearest-neighbor graphs were constructed for clustering. Next, Uniform Manifold Approximation and Projection (UMAP) embeddings were generated with n.neighbors = 30, min.dist = 0.3, and cosine distance. (Becht et al. 2018). For all analyses, clustering was evaluated across resolutions 0.1 to 1.0 in increments of 0.1, and the final resolution was selected from this range, generally using the resolution that gave the minimum number of clusters to resolve known cell types. Unless otherwise noted, clustering used the Louvain algorithm with multilevel refinement (Blondel et al. 2008). Cell types were annotated by performing cluster marker analysis with Seurat::FindAllMarkers, and cluster identities were assigned based on the expression of curated marker genes (Brann et al. 2020; Gadye et al. 2017) (**Table S1**). Analysis details specific to each cell subset are below.

#### Whole OE analysis

For the 82,677-cell whole OE Seurat object, UMAP embedding was performed using the first 16 PCs, and unsupervised SNN graph clustering using the original Louvain algorithm at resolution 0.3 revealed 21 clusters (**Fig. S2**). Clusters corresponded to cell types of the sensory OE and surrounding tissue. Respiratory epithelial cell types included respiratory basal cells (cluster 6: *Krt17*^+^, *Reg3g*^+^) and one cluster expressing secretory, respiratory, BG, and goblet cell markers (cluster 15: *Tff2*^+^, *Muc5b*^+^, *Reg3g*^+^, Wfdc18^+^). To differentiate between respiratory goblet cells and cells of the Bowman’s gland in this cluster, we reclustered cells at a resolution of 1.0 and differentiated the cell types by expression of *Sox9* and *Wfdc18* (Bowman’s glands) or *Tff2* and *Muc5b* (respiratory goblet cells) (**Fig. S2**) (Montoro et al. 2018). We also identified olfactory bulb-derived cells (cluster 18: *Dcc*^+^, *Pou3f2*^+^; cluster 5: *Igfbpl1*^+^, *Dlx2*^+^), olfactory ensheathing cells (cluster 17: *Npy*^+^, *S100b*^+^), odontogenic epithelium (cluster 19: *Amelx*^+^, *Enam*^+^), fibroblasts (cluster 10: *Col1a1*^+^, *Col1a2*^+^), two immune-enriched myeloid clusters, including a macrophage-like cluster (cluster 11: *C1qb*^+^, *Cd68*^+^) and an inflammatory monocyte-like cluster (cluster 14: *S100a8*^+^, *S100a9*^+^, *Il1b*^+^). A cluster with mixed cell-type gene expression was also identified and may represent doublets (cluster 20: *Trp63*^+^, *Krt15*^+^, *Omp*^+^, *Gap43*^+^).

Remaining clusters consisted of cell types in the main sensory OE lineages: HBCs (cluster 9: *Trp63*^+^, *Krt5*^+^), GBCs (cluster 7: *Ascl1*^+^, *Kit*^+^), INPs (clusters 4, 13: *Neurod1*^+^, *Neurog1*^+^), iOSNs (clusters 1, 3: *Ebf2*^+^, *Gap43*^+^), mOSNs (clusters 0, 2, 12: *Omp*^+^), sustentacular cells (cluster 8: *Cyp2a5*^+^, *Il33*^+^), microvillus cells (cluster 16: *Ascl3*^+^, *Foxi1*^+^), and cells of the Bowman’s gland (the retained subset of cluster 15: *Sox9*^+^, *Wfdc18*^+^).

### Sensory OE analysis

Next, we removed all cells not in the main sensory OE lineages from the Seurat object. Cells were reprocessed as described above using 14 PCs for downstream analysis. Clustering at resolution 0.4 yielded 14 clusters including a cluster of potential doublets (cluster 12) that expressed genes consistent with multiple lineages, including high expression of *Wfdc18*, *Sox9*, *Muc5b*, *Reg3g*, and *Cftr*, together with epithelial markers such as *Krt8*, *Krt18*, and *Krt23*. After removal of cluster 12, 62,190 cells remained (**Fig. S3A,B**).

This object was then reclustered using 12 PCs at resolution 0.6 and annotated based on gene expression (**Table S1**). This yielded 14 clusters representing the known major OE cell types: mature olfactory sensory neurons (mOSN 1, mOSN 2, mOSN 3: *Omp*^+^), immature olfactory sensory neurons (iOSN1, iOSN2, iOSN3: *Gap43*^+^*, Ebf2*^+^), early and late immediate neuronal progenitors (INPs: *Neurod1*^+^, *Neurog1*^+^, *Ebf1*^+^, and *Ebf2*^+^; early INPs show higher expression of the proliferation genes *Mki67* and *Top2a*), Sus cells (*Cyp2g1*^+^, *Cyp1a2*^+^, *Il33*^+^, *Cyp2a5*^+^, *Sec14l3*^+^), HBCs (*Krt5*^+^, *Trp63*^+^, *Krt17*^+^, *Nrcam*^+^), GBCs (*Kit*^+^, *Ascl1*^+^, *Mki67*^+^), BG cells (*Sox9*^+^*, Wfdc18*^+^), and MV/tuft MV cells (*Trpm5*^+^, *Ascl3*^+^, *Foxi1*^+^). Breakdown of cluster membership by individual animal rather than age is available in **Fig. S3.** All ages were represented in all sensory OE clusters, and cluster distribution was consistent among animals in the same age group (**Fig. 2B; Fig. S3C**).

### Ascl1^+^ subset analysis

To isolate *Ascl1*^+^ cells for further characterization (**Fig. 3**, **Fig. 4C, D; Table S3**), 4,478 *Ascl1*^+^ (*Ascl1* expression > 0) cells were extracted from the lineage object, followed by re-processing as described above using 16 PCs for downstream analysis. Clustering at resolution 0.7 yielded 12 clusters, including 5 clusters consistent with the neuronal lineage (expressing *Ebf2* and other neuronal markers), which were removed (**Fig.S4**), leaving 2,732 cells in 7 clusters. For **Fig. 3A**, module scores were calculated using Seurat::AddModuleScore with ctrl = 5. Gene sets used to specify modules were: GBC (*Kit, Ascl1, Mki67*), sustentacular (*Cyp2g1, Cyp2a5, Sec14l3, Il33*), HBC (*Trp63, Krt5, Krt14, Krt17, Nrcam*), microvillus (*Cftr, Foxi1, Ascl3, Pou2f3*), and Bowman’s gland (*Sox9, Wfdc18*).

For **Fig. 4**, differential marker analysis was performed with Seurat::FindAllMarkers. We then subset the identified genes to only those included in a previously published list of 1,639 mouse transcription factors (Van den Berge et al. 2026). The top markers for each cluster were ranked by adjusted p-value. If a gene appeared among the top-ranked markers of multiple clusters, it was assigned only once, to the cluster in which its average expression was highest.

### HBC subset analysis

HBCs were isolated for further characterization (**Fig. 6; Table S3**) by subsetting the HBC cluster (cluster 9 of the olfactory lineage Seurat object at resolution 0.6), followed by reprocessing of the data as described above using 14 PCs. Clustering at resolution 0.3 yielded 6 clusters.

Cell-cycle phase was assigned using Seurat::CellCycleScoring, which produces module scores for cell cycle based on canonical marker genes and assigns each cell to its most likely place in the cell cycle. We first converted the human genes provided into mouse genes by altering their capitalization (ex: “PCNA” to “Pcna”) then calculated cell cycle scores to predict the state of each cell.

#### Trajectory inference

Trajectory inference was performed using *slingshot* (Street et al. 2018). We first subset the Seurat object to exclude tuft MV and Bowman’s gland cells, both of which were relatively rare populations (the two clusters together represented 1.37% of total cells), leaving 61,337 cells. Plotting marker genes revealed a group of cells within the Sus cluster with high levels of expression of neuronal genes. Clustering at resolution 4.0 restricted these cells to one neuronal-like cluster of Sus cells (394 cells; **Fig. S5; Table S4**), which was removed. We then reprocessed the data using 18 PCs to produce the final object, which contained 60,265 cells expressing 32,285 genes. Cells retained their original cell type designation as assigned in the whole OE object (12 clusters). *Slingshot* was performed using the first 5 principal components, specifying the *Ascl1*-high cluster as the starting cluster and HBCs, Sus cells and MV cells as endpoints. This identified four lineages: mOSN, HBC, Sus, and MV.

### Lineage-specific gene identification

Differential expression along pseudotime was performed using tradeSeq (Van den Berge et al. 2020) with knots set to 6. We limited the analysis to a gene set consisting of the union of the 2,000 most highly variable genes in the Slingshot object, excluding olfactory receptor genes, and a list of 1,639 mouse transcription factors (Van den Berge et al. 2026).

To identify genes that changed over the course of each lineage, we applied tradeSeq::startVsEndTest to the fitted generalized additive models. This test compares the fitted expression at the beginning and end of each lineage and returns a lineage-specific Wald statistic and p-value for each gene. For each lineage, p-values were adjusted using the Benjamini-Hochberg procedure to control the false discovery rate (FDR).

Genes were considered upregulated along a lineage if they showed a significant start-to-end increase (at nominal FDR level of 0.05) and a positive fitted log-fold change. To define stringent lineage-specific genes, we further required that a gene reach its highest fitted endpoint expression in that lineage and that and that the fitted endpoint expression margin versus the next highest lineage was > 1. Thus, lineage-specific genes were defined as genes that both increased significantly along a lineage and ended higher in that lineage relative to the other branches.

### CellChat analysis

For niche signaling analysis, we used CellChat v2.2.0 (Jin et al. 2021) on the whole OE dataset. Clusters were defined based on cluster assignments from the whole OE object, and consisted of fibroblasts (cluster 10), GBCs (cluster 7), HBCs (cluster 9), inflammatory monocyte-like cells (cluster 14), INPs (clusters 4 and 13), iOSNs (clusters 1 and 3), macrophage-like cells (cluster 11), microvillus cells (cluster 16), mOSNs (clusters 0, 2 and 12), olfactory ensheathing cells (cluster 17), respiratory basal cells (cluster 6), olfactory bulb-derived cells (clusters 5 and 18), sustentacular cells (cluster 8), cells of the Bowman’s gland (cluster 15 cells where RNA_snn_res.1 == 31: cluster 15a), respiratory goblet cells (remaining cluster 15 cells: cluster 15b), and odontogenic epithelium (cluster 19). Cluster 20 was manually removed prior to analysis.

We excluded classes from the CellChat analysis if at least one age contained fewer than 10 cells, which removed Bowman’s glands, odontogenic epithelium, olfactory ensheathing cells, and olfactory bulb clusters from consideration.

The annotated Seurat object was split by developmental age, and CellChat analysis was performed separately for each age. For each age, the CellChat object was created from the Seurat RNA assay using CellChat::createCellChat with cell-type annotations as the grouping variable and the mouse ligand-receptor database (CellChatDB.mouse). Standard CellChat preprocessing was applied using CellChat::subsetData, CellChat::identifyOverExpressedGenes, and CellChat::identifyOverExpressedInteractions, followed by inference of communication probabilities with CellChat::computeCommunProb(population.size = TRUE) in order to account for for unequal population sizes of senders and receivers. Inferred interactions were further filtered with CellChat::filterCommunication(min.cells = 10) and pathway-level communication probabilities were computed with CellChat::computeCommunProbPathway, and aggregated with CellChat::aggregateNet.

Age-specific CellChat objects were then merged using CellChat::mergeCellChat for downstream comparison across ages. To quantify incoming signaling to GBCs, communication tables were extracted using CellChat::subsetCommunication and filtered for target == GBC. Overall incoming signal strength was calculated from the summed communication probabilities across incoming ligand-receptor interactions, and sender-specific incoming strength was calculated by summing probabilities across all interactions from each sender cell type to GBCs within each age. Pathway level summaries were calculated using the netP communication table after filtering for GBC as receiver. These values were then used for cross-age comparisons and for pathway level summaries (**Fig. 7A, Fig. S6, Table S5**).

To analyze total incoming signaling at P5 vs P22 (**Fig. 7A,B; Fig. S6**), we extracted and summed incoming CellChat communication probabilities from each of the 12 retained sender cell types to GBCs at P5 and at P22. We then calculated P5 - P22 for each sender-pathway or sender-ligand-receptor group. Positive values indicate stronger predicted signaling at P5 while negative values indicate stronger signaling at P22. For figures, sender cell types were restricted to the top 6 senders with the largest summed incoming signal to GBCs at P5, ranked by summed ligand-receptor communication probabilities from the CellChat net table. In **Fig. 7B**, incoming pathway probabilities were extracted from the CellChat netP slot, summed within each sender and age, and compared between P5 and P22. The top 8 pathways per selected sender, ranked within each sender by absolute P5 - P22 delta, are displayed.

## Data and code availability

Raw images and primary data files supporting this study are available from the corresponding author upon request. Sequencing data has been deposited to GEO under the accession number GSE341251. Custom code is available from the corresponding author upon request and will be made publicly available in a version-controlled repository.

## List of Supplementary Info

Supplementary Figures (Figs. S1-S6)

Supplementary Table S1. WholeOE QC / annotation / lineage selection

Supplementary Table S2. Histology data and statistics

Supplementary Table S3. Markers / Ascl1_pos programs / HBC subsets

Supplementary Table S4. Trajectory / tradeSeq analysis

Supplementary Table S5. CellChat analysis

Supplementary Table S6. Key resources / antibodies / software / computational environment

## Supplemental Figures

**Figure S1:**
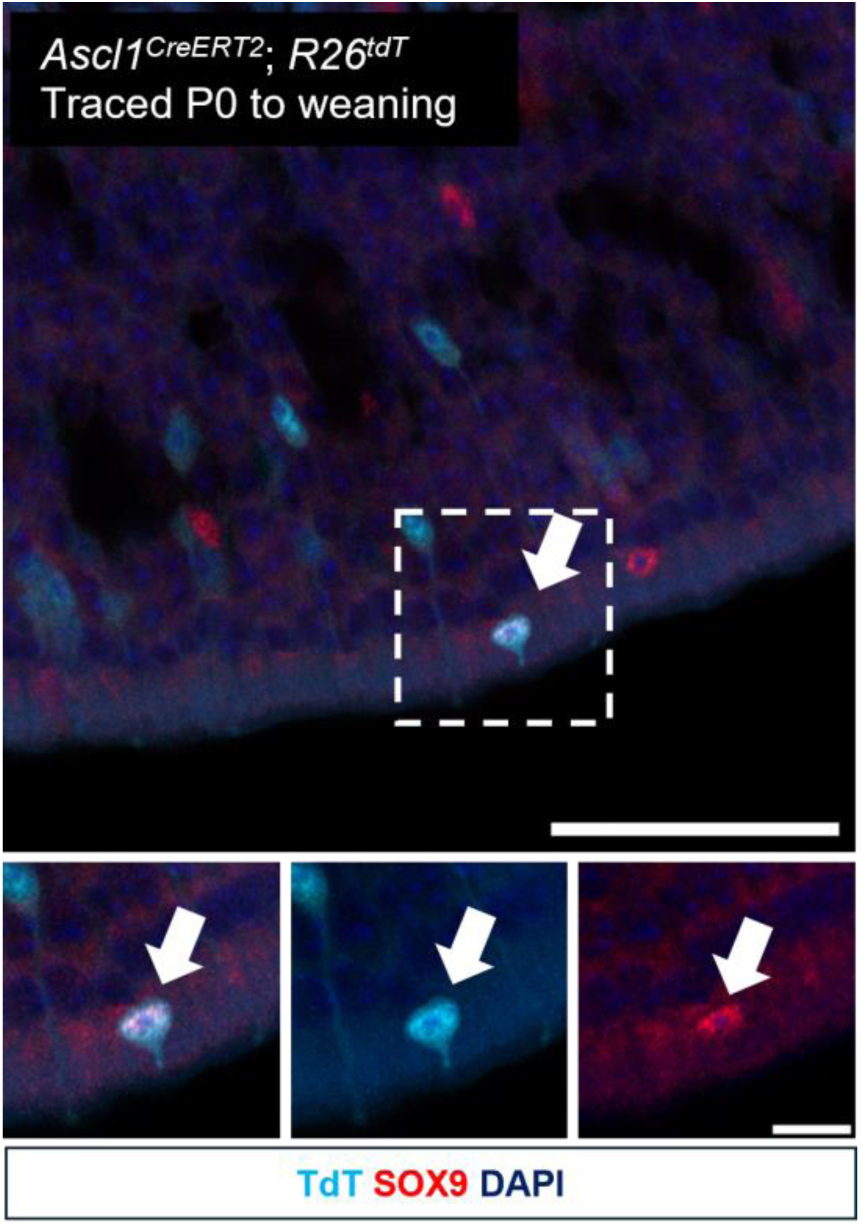
Microvillus cell labeled by *Ascl1^CreERT2+^* lineage tracing. An example of a rare tdT^+^ (cyan) microvillus cell (SOX9^+^; magenta) traced from P0 and visualized at weaning. Arrow and boxed region indicate the traced cell; lower panels show higher-magnification views of the boxed region. Scale bar, top panel: 50 μm; inset panels: 10 μm.

**Figure S2:**
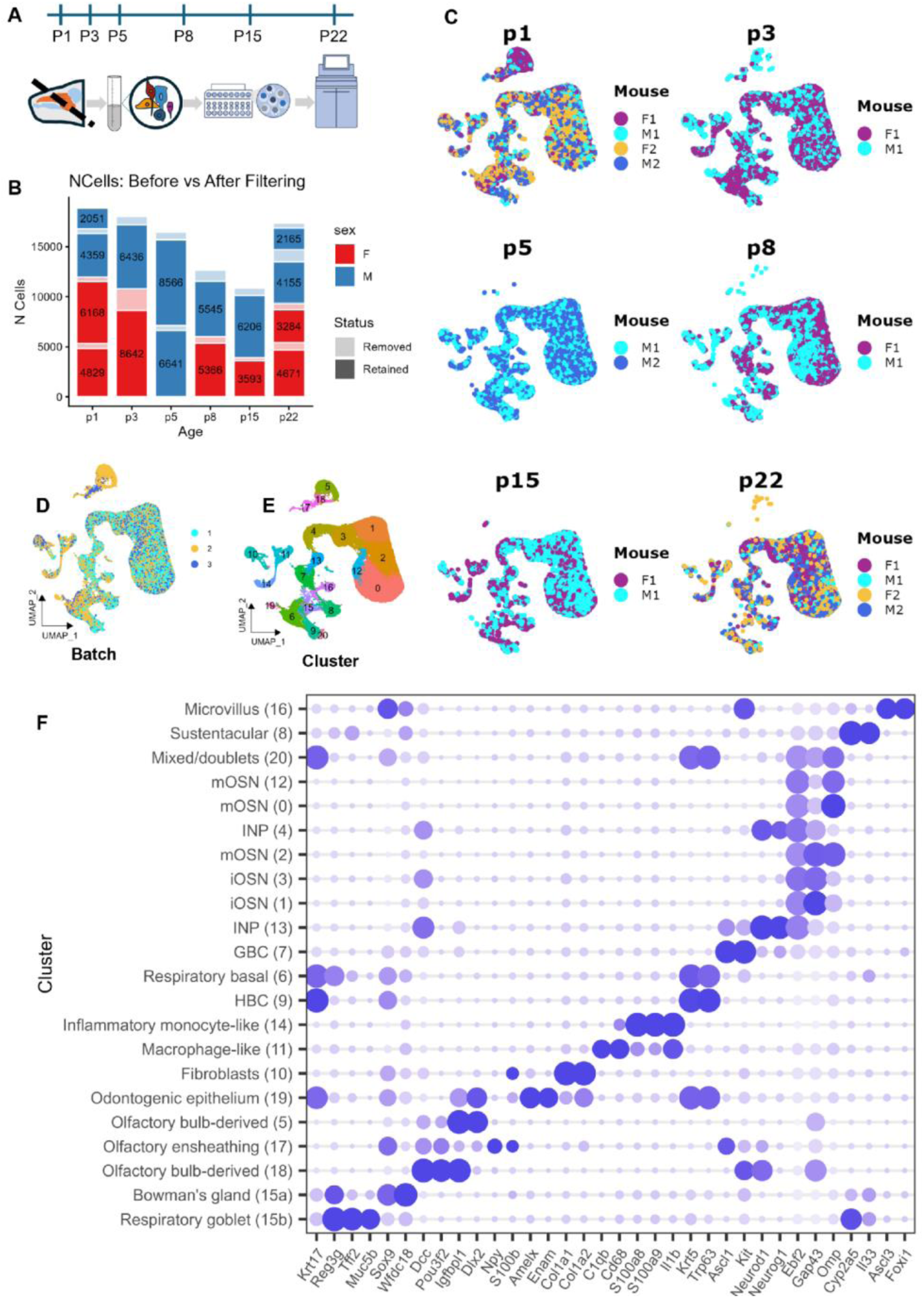
Single-cell RNA-seq processing, quality control, and clustering overview. **(A)** Overview of single cell processing pipeline. **(B)** Metadata indicating ages and sexes of removed cells after QC filtering. **(C)** UMAPs colored by animal, split by age show good inter-sample mixing, with no clear separation by animal. **(D)** UMAP colored by dissection batch. Batch 2 (yellow), containing P1 animals, has relatively more respiratory epithelium and olfactory bulb than other batches due to the difficulty of dissecting these small animals. **(E)** UMAP colored by cluster. **(F)** Dotplot of marker genes by cluster.

**Figure S3:**
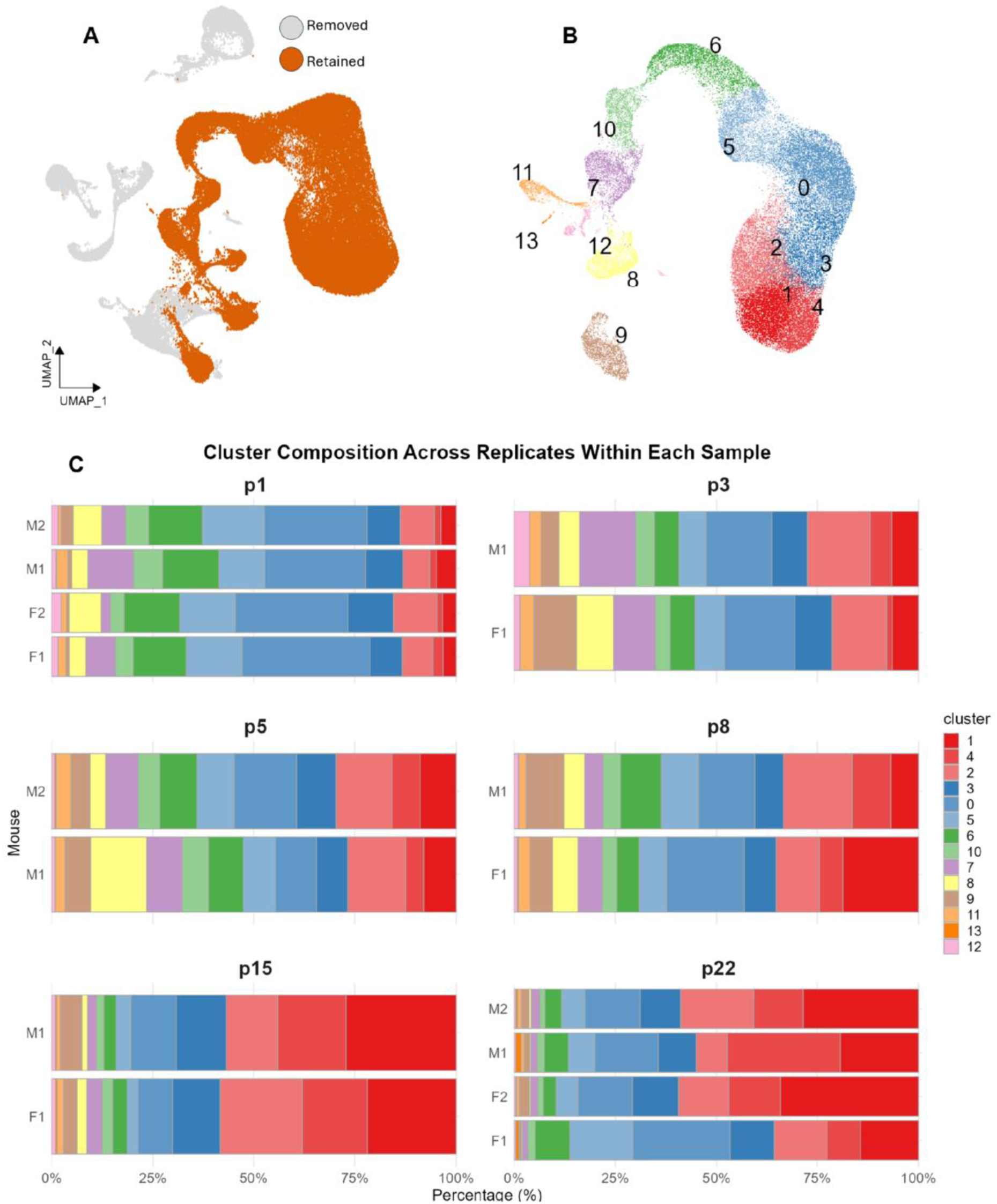
Sensory OE cell types by sample. **(A)** UMAP of whole OE showing cells removed (grey, non-sensory) and retained (orange, sensory) for downstream processing. **(B)** UMAP of sensory OE labeled by cluster. **(C)** Stacked bar plots showing the percentage of cells from each animal in each cluster.

**Figure S4:**
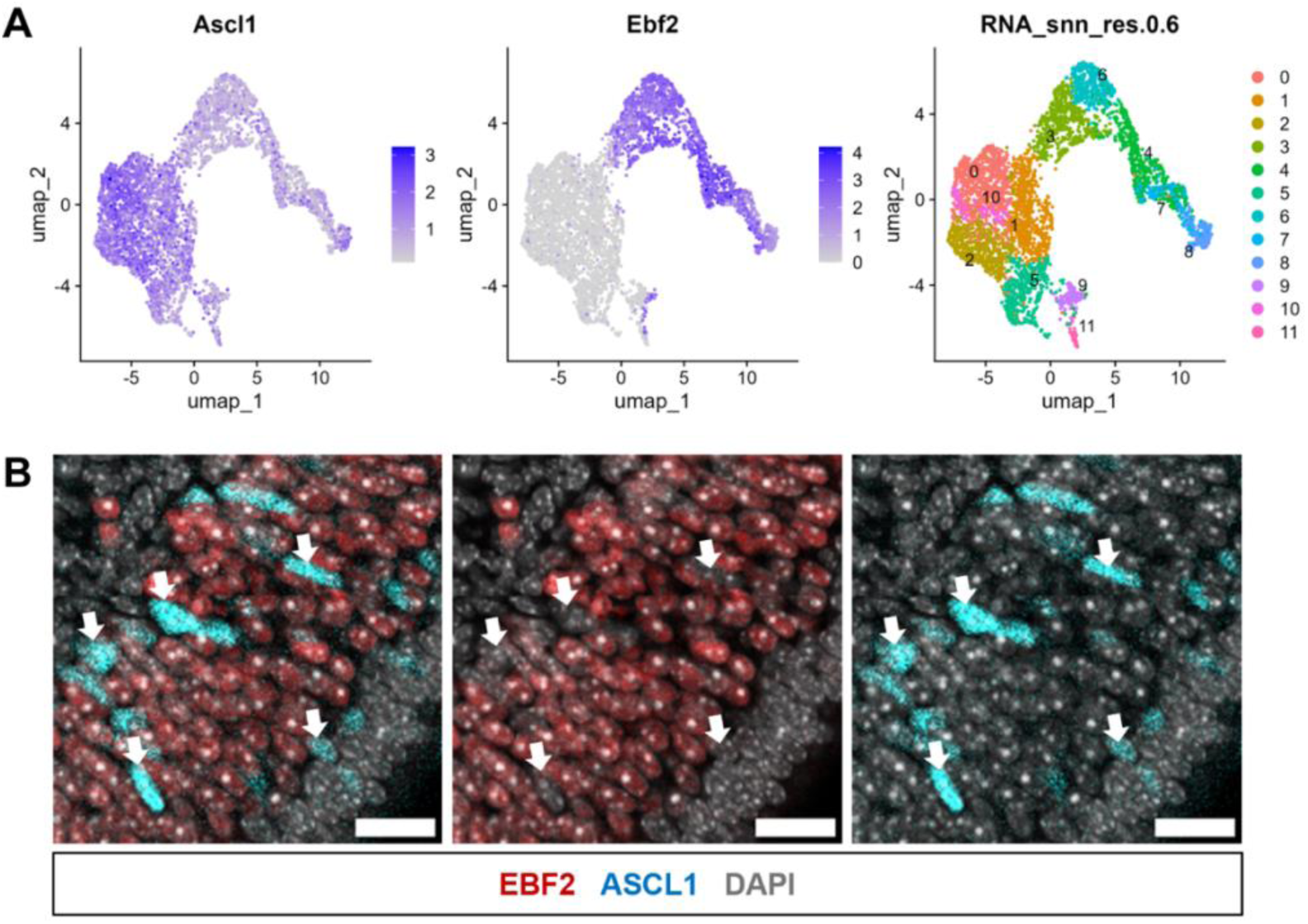
ASCL1 and EBF2 protein are not co-expressed at P1. (**A)** UMAPs of *Ascl1*^+^ cell subset colored by *Ascl1* expression (left), *Ebf2* expression (middle), or cluster membership (right). **(B)** Representative immunofluorescence images of wild-type C57BL/6 OE at P1 showing EBF2 (red), ASCL1 (cyan), and DAPI (gray); Arrows indicate cells expressing ASCL1 but not EBF2. Scale bar: 20 μm

**Figure S5:**
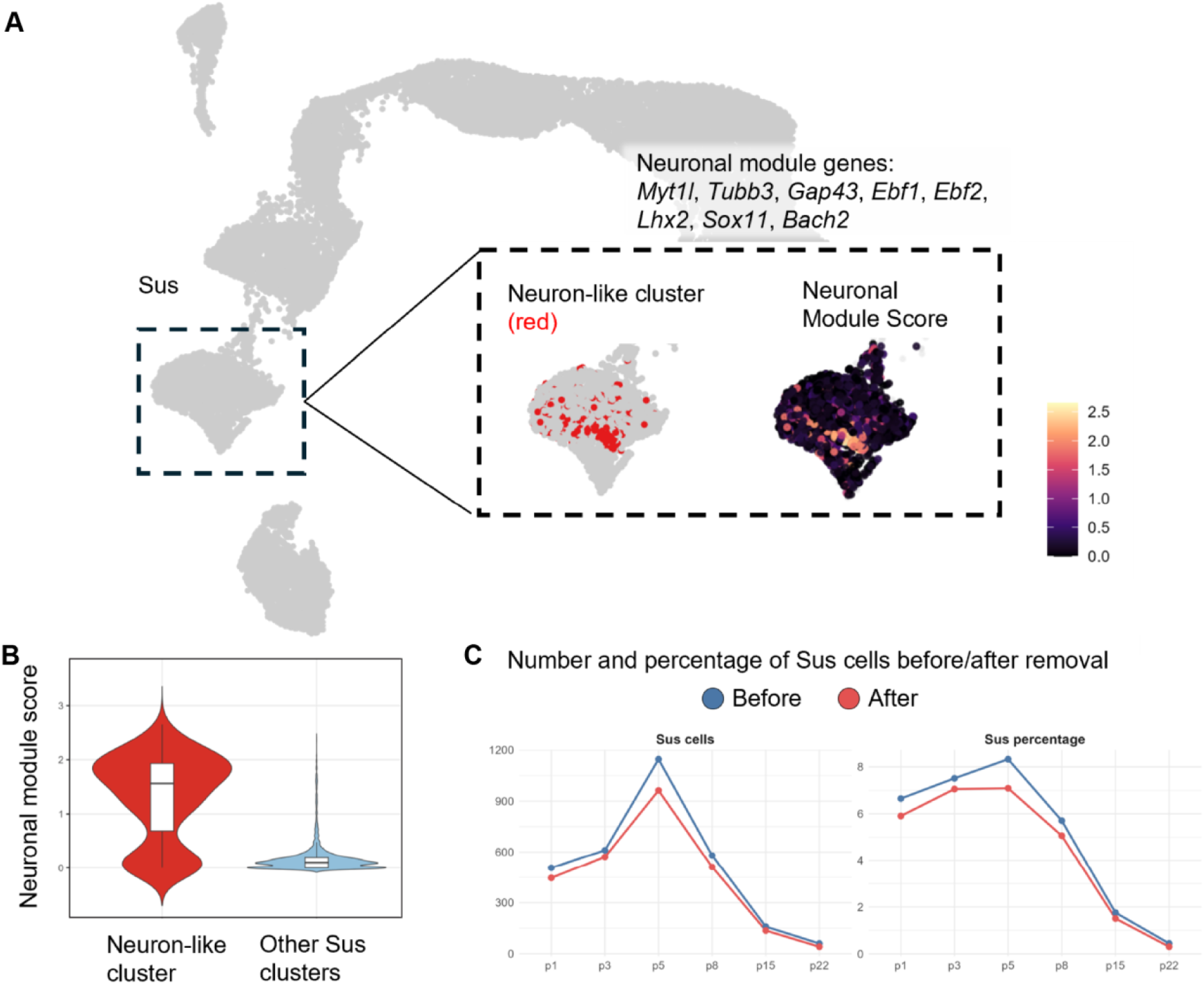
Identification and removal of a neuronal-gene expressing sustentacular cell cluster. **(A)** UMAP of sensory OE cells highlighting the sustentacular (Sus) compartment (dashed box). **(B)** Sus cells with cluster 57 highlighted in red (left) and the corresponding neuronal module score overlaid (right), showing enrichment of neuronal gene expression within cluster 57. The neuronal module was calculated from the genes *Myt1l*, *Tubb3*, *Gap43*, *Ebf1*, *Ebf2*, *Lhx2*, *Sox11*, and *Bach2*. **(C)** Violin plot comparing neuronal module scores in cluster 57 versus all other Sus clusters. **(D)** Number and percentage of Sus cells before and after removal of cluster 57 across developmental time points.

**Figure S6:**
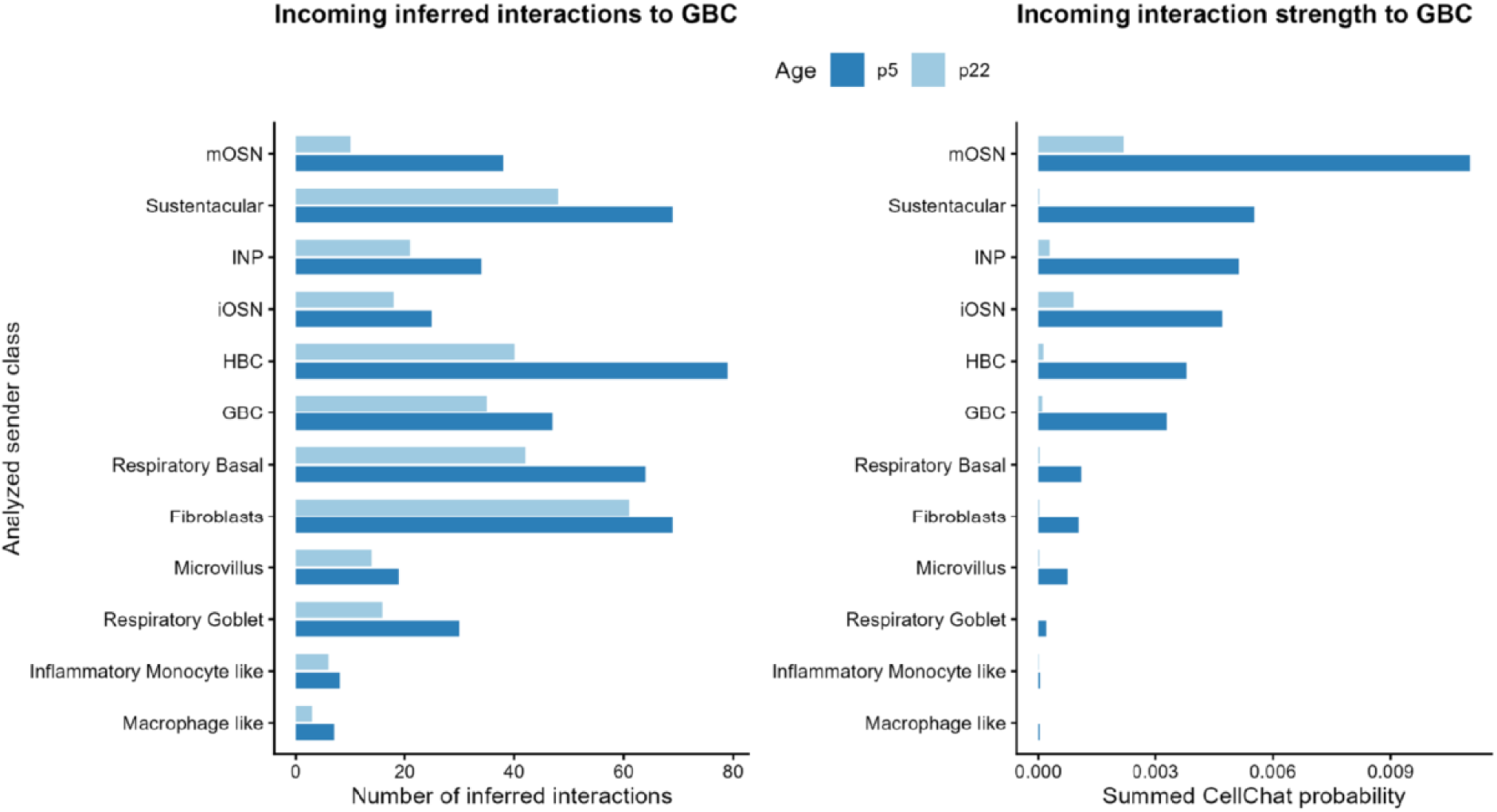
Summary of incoming signals to GBCs from all sender cell types considered. (Left) Number of inferred incoming interactions to GBC from senders; (Right) summed CellChat interaction strength from each sender class. Predicted incoming signaling was strongest from mOSNs, Sus cells, INPs, iOSNs, HBCs, and GBCs, whereas respiratory, immune, MV, and fibroblast inputs contributed comparatively weak signals.

